# Protein-stabilizing and neurotransmission-potentiating activities of a synaptic chaperone modify spinal muscular atrophy in model mice

**DOI:** 10.64898/2026.02.23.707472

**Authors:** Yoon-Ra Her, Andrea Fuentes-Moliz, Rashmi Kothary, Lucia Tabares, Umrao R. Monani

## Abstract

Spinal muscular atrophy (SMA) is an oft-fatal infantile-onset neuromuscular disease caused by low SMN protein. Administration of SMN-inducing agents to SMA newborns prevents early mortality, but therapeutic outcomes vary considerably, and disease mechanisms remain poorly understood. Genetic modifiers can provide clues to disease mechanisms and serve as targets for novel treatments. Here, we describe how one such modifier suppresses SMA in model mice. We show that the modifier, an Hspa8^G470R^ synaptic chaperone variant we previously identified, functions beyond an already defined role as an *SMN2* splice-switcher. Even in mice lacking the *SMN2* gene, the modifier, whether expressed genetically or exogenously, potently suppressed disease, preventing motor neuron degeneration, ameliorating neuromuscular dysfunction and extending lifespan more than ten-fold. Unexpectedly, this was once again associated with incremental SMN increase – an outcome we discovered is linked to Hspa8^G470R^-mediated autophagy, effects of the modifier on autophagy-associated intermediate complexes and, ultimately, reduced SMN turnover. Interestingly, however, Hspa8^G470R^ also stimulated neuromuscular transmission significantly, raising the effective, functional readily releasable pool of motor neuronal synaptic vesicles. This effect was not limited to mutants alone but apparent in healthy controls too and did not correlate with mere increase in SMN. Combined, these outcomes suggest that Hspa8 governs neuromuscular function in several ways including direct effects on synapses. Mechanisms revealed here shed additional light on pathways gone awry in SMA – ones that might be modulated to develop or refine therapies for neuromuscular disorders at large.

## Introduction

Homozygous mutations in the Survival Motor Neuron 1 *(SMN1)* gene result in low levels of its namesake SMN protein and trigger the frequently fatal pediatric-onset neuromuscular disease, spinal muscular atrophy^1^. Several SMN-inducing agents are now approved for the treatment of SMA^2^ and newborn screening for the disease routinely offered to new parents^3^. These developments have radically altered disease outcomes, particularly when infants are treated soon after birth. Still, none of the SMA therapies is curative, with many infants, especially those with 2 *SMN2* copies – predictive of the severest form of the disease – failing to meet developmental motor milestones, notwithstanding early treatment^4^. Therapeutic intervention in these patients has also unmasked previously unappreciated phenotypes, a consequence of having prevented the early mortality associated with severe SMA^5,6^. These outcomes may be explained by insufficient SMN induction following postnatal therapy and/or critical requirements for the protein *in utero*. However, they are also impacted by an incomplete understanding of the most critical SMN-linked pathways that are perturbed in SMA and that must be normalized to prevent or reverse disease.

Genetic modifiers of disease constitute important windows into the selective vulnerability of organ systems to deficiencies in essential proteins like SMN. Indeed, identifying SMA modifiers has been informative, linking SMN to neuromuscular junction (NMJ) organizers such as Agrin, MuSK and Dok7, to perturbed neuronal endocytosis and to factors essential for neurotransmission, e.g., Syt2 (Synaptotagmin2) and the P/Q type Ca^2+^ channels in motor neurons^7–13^. Still, exactly how SMN alters the expression or localization of these proteins is unclear. Consequently, the extent to which pathways associated with these proteins influence the SMA phenotype remains to be fully explained. We previously found that a G470R variant of the synaptic Hspa8 chaperone protein potently suppresses SMA in model mice^14^. Biochemical studies revealed that the modifier functions as an *SMN2* splice modulator, modestly raising functional SMN protein. However, we also found that it enhanced synaptic SNARE complex levels significantly, an observation consistent with known Hspa8 functions^15,16^. This raised the prospect of a *direct* protective effect of Hspa8^G470R^ on SMA NMJs but one confounded by the co-incidental *SMN2* splice-switching properties of the modifier; increased SMN protein resulting from higher full-length (FL) SMN transcript in the presence of Hspa8^G470R^ may be argued to be the exclusive driver of enhanced neurotransmission and healthier NMJs in modified mutant mice. Here, we attempted to dissociate any direct ameliorative effect of Hspa8^G470R^ from one mediated by its role as an *SMN2* splicing modulator by investigating if model mice *devoid* of the copy gene nevertheless derive benefit from the modifier. We found that Hspa8^G470R^ is equally potent in suppressing disease in such mutants, preserving spinal motor neurons, mitigating neuromuscular pathology and extending lifespan significantly. Intriguingly, however, the modifier once again raised SMN – by stabilizing the protein, via its effects as a cellular chaperone of clients destined for autophagic degradation. This effect was nevertheless accompanied by enhanced NMJ neurotransmission and, interestingly, a marked increase in the readily releasable pool (RRP) of synaptic vesicles that exceeded the RRP of even healthy controls – notwithstanding the lower SMN in modified mutants. Our results not only assign a new mechanism of action to the Hspa8^G470R^ modifier in regulating SMN protein but also bolster the notion that it equally likely acts directly at synapses to potentiate neurotransmission and suppress disease. The multiple mechanisms of action we attribute to this variant in modulating the SMA phenotype make it and the pathways it affects appealing targets not just to refine treatments for this pediatric disorder but also to mitigate allied motor neuron conditions.

## Results

### The Hspa8^G470R^ variant suppresses overt disease and neuromuscular pathology in the Smn^2B/-^ mouse model of SMA

We previously employed ‘SMNΔ7’ model mice^17^ harboring the human *SMN2* gene to map and identify a potent suppressor of the SMA phenotype^14^. The Hspa8^G470R^ disease suppressor raised SMN levels by functioning as an *SMN2* splice-modulator. However, it also potentiated NMJ neurotransmission and stimulated synaptic SNARE complex assembly, suggesting a second, SMN-independent disease-modifying effect. Recognizing the challenge of discerning the relative contribution of each of these effects to disease rescue in *SMN2*-expressing mutants, we sought to examine modifier-driven outcomes in model mice that do not rely on the human copy gene. For this, we turned to the *Smn^2B/-^* SMA mutant, wherein the murine *Smn* allele is engineered to mis-splice and produce low SMN^18,19^. Because strain background influences disease outcomes, and considering that our earlier study was conducted on mutants derived on the FVB/N background, we made certain that our founder *Smn^2B^* breeders were also FVB/N. Single polymorphism nucleotide (SNP) analysis using 146 markers informative for the FVB/N and C57BL/6J strain confirmed that the breeders derived 99.31% of their genomes from the FVB/N strain (Table S1). We thus proceeded with these animals and bred them to Hspa8^G470R^ mice on the same strain background to generate mutants for our study. We found that disease in *Smn^2B/-^* mutants bearing Hspa8^G470R^ was markedly suppressed. Median survival of mutants with one Hspa8^G470R^ allele increased from PND19 to PND238; mutants homozygous for the modifier survived even longer, indicative of dose dependency (Fig. 1A). Additionally, overall health as assessed by weight gain during the first three postnatal weeks of life improved (Fig. 1B and Fig. S1A) and agility of *Smn^2B/-^;Hspa8^G470R^* mutants between PND8 and PND10, as determined by righting ability, was significantly enhanced relative to that of mutants absent the modifier (Fig. 1C). Finally, we also found that SMA mutants bearing the modifier displayed greater grip strength than cohorts without it (Fig. 1D).

**Figure 1.**
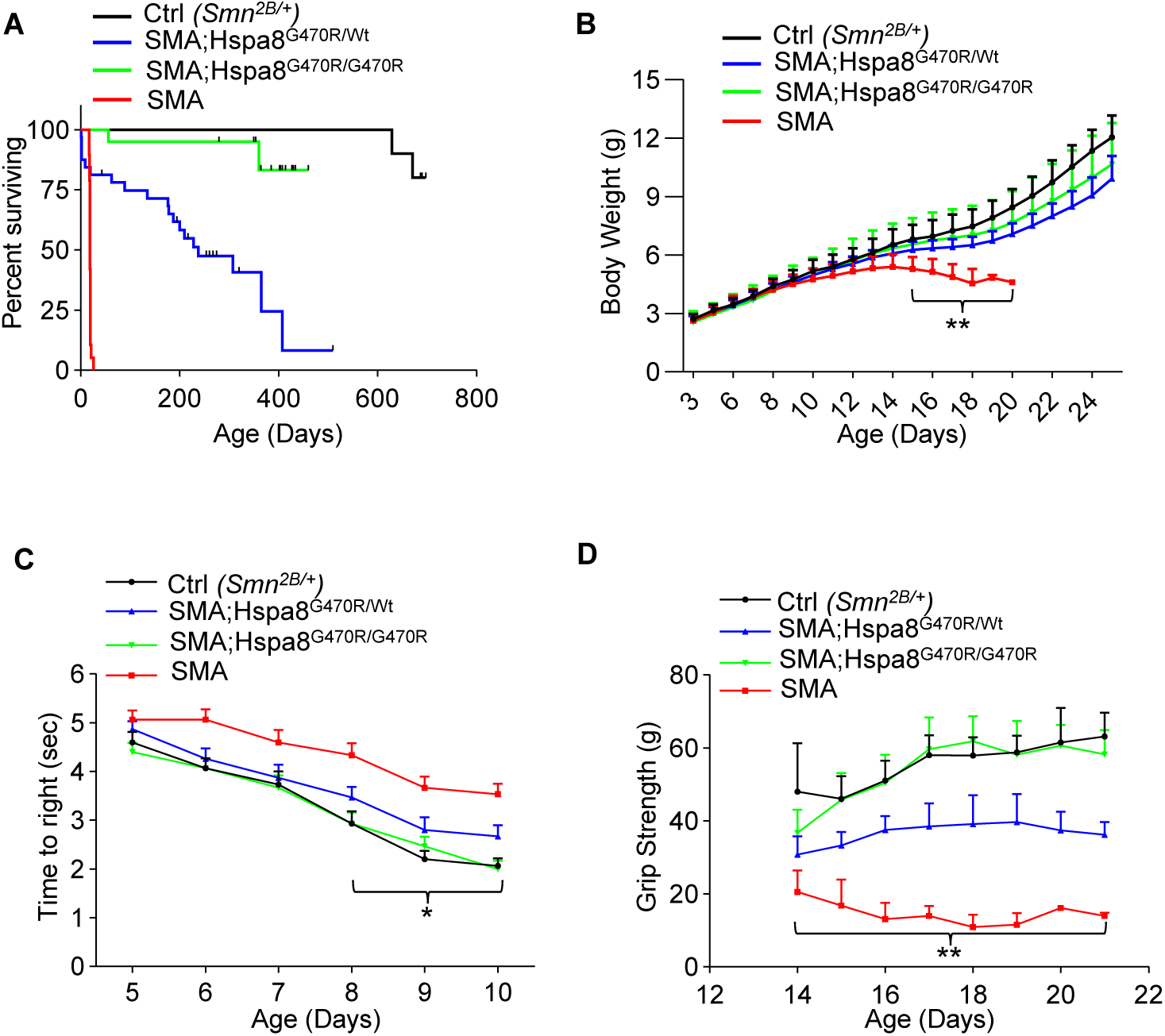
The Hspa8^G470R^ variant mitigates overt disease in SMA model mice. **(A)** Kaplan-Meier survival curves depicting significant enhancement of mutant lifespan in the presence of the Hspa8^G470R^ modifier. *P* < 0.0001 between SMA and SMA;Hspa8^G470R^ mutants, log-rank test, n = 10 – 32 mice of each cohort. **(B)** Mutants continue to gain weight in the presence of the modifier. **, *P* < 0.01, between SMA and SMA;Hspa8^G470R^ mutants, one-way ANOVA, n = 14 – 22 mice. **(C)** Quantified results of righting time depicts improved motor performance of mutants expressing Hspa8^G470R^. **, *P* < 0.05, between SMA and SMA;Hspa8^G470R^ mutants, two-way ANOVA, n = 15 mice of each genotype. **(D)** Graph showing that Hspa8^G470R^ restores grip strength to SMA mutants. Note: **, *P* < 0.01 between SMA and SMA;Hspa8^G470R^ mutants; two-way ANOVA, n = 3 – 19 mice. Data: mean ± SEM

Neuromuscular pathology is a signature feature of SMA and is expected to be alleviated by disease suppressors. Accordingly, we began by examining muscle tissue of PND15 *Smn^2B/-^* mutants with and without the modifier. We found that myofibers in both the proximal triceps muscles as well as distal gastrocnemius of mutants harboring Hspa8^G470R^ were significantly larger than those of mutants without the modifier (Fig. 2A, B and Fig. S1B), and this was reflected in frequency histograms revealing a shift toward fibers with greater cross-sectional areas in modified mutants (Fig. S1C, D). We next quantified spinal motor neurons in our various cohorts of mice. Expectedly, morphometric counts revealed fewer (∼40%) cell bodies in lumbar spinal cord of *Smn^2B/-^* mutants relative to healthy controls. In contrast, we found no evidence of significant motor neuron loss in mutants expressing Hspa8^G470R^ (Fig. 2C, D). To complete our assessment of neuromuscular health, we conducted a detailed anatomical assessment of the NMJs in the PND15 mutants and healthy controls. We found that NMJ abnormalities in *Smn^2B/-^* were either abolished or significantly mitigated by the modifier. Thus, for instance, endplate denervation in muscles of *Smn^2B/-^;Hspa8^G470R/Wt^* mutants, heterozygous for the modifier, was significantly reduced but not to control levels; NMJ innervation in mutants homozygous for Hspa8^G470R^ was further improved and equivalent to that of controls (Fig. 2E, F and Figs. S1E). We observed similar patterns of rescue when NMJs were examined for nerve terminals abnormally swollen with neurofilament (NF) protein and when endplate size and complexity – as assessed by perforations – was determined (Fig. 2G – I).

**Figure 2.**
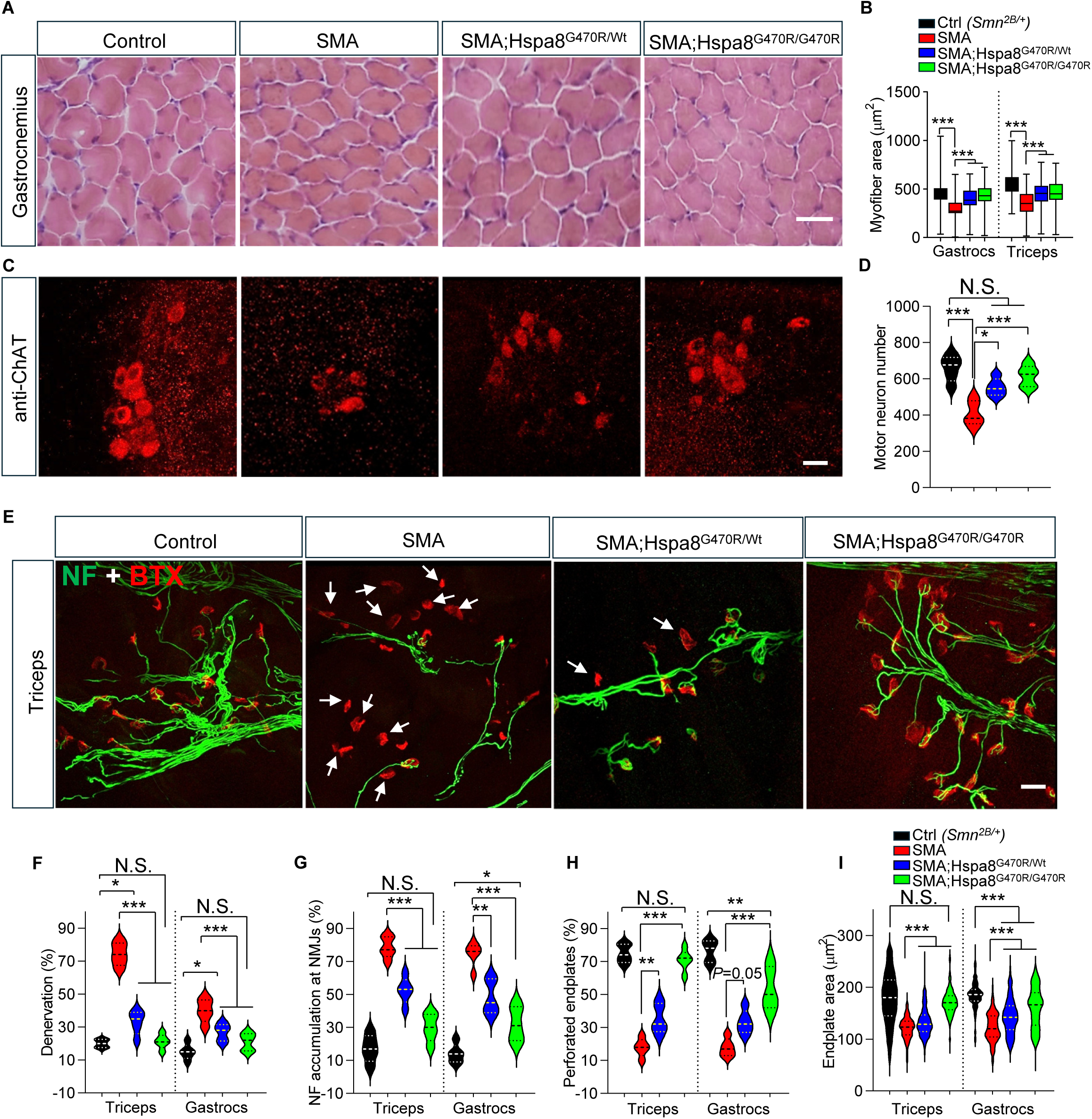
The Hspa8^G470R^ variant lessens neuromuscular pathology in SMA model mice. **(A)** Representative H&E-stained sections of gastrocnemius muscles from PND15 controls and mutants with and without the Hspa8^G470R^ modifier. Scale bar – 25μm. **(B)** Quantified myofiber sizes in controls and SMA mutants with or without Hspa8^G470R^. ***, *P* < 0.001, Kruskal-Wallis test, n > 1000 fibers from N = 3 mice of each cohort. **(C)** Representative lumbar spinal cord sections from the various mouse cohorts depicting ventral horn cells. Scale bar – 20μm. **(D)** Morphometric counts of ventral horn cells in controls and SMA mutants with or without Hspa8^G470R^. *, ***, *P* < 0.05 and *P* < 0.001, respectively one-way ANOVA, n = 5 mice of each genotype. **(E)** Immunostains of NMJs in muscle from controls and mutants bearing or devoid of the Hspa8^G470R^ modifier. Arrows denote denervated endplates. Scale bar – 20μm. Quantification of **(F)** denervated endplates, **(G)** endplates exhibiting abnormal swellings of NF protein in nerve terminals, **(H)** endplate complexity and **(I)** endplate size, in controls and mutants with or without the SMA modifier. Note: *, **, ***, *P* < 0.05, *P* < 0.01 and *P* < 0.001 respectively, one-way ANOVA, n > 100 endplates from N = 3 – 5 mice of each cohort (panels F – H); Kruskal-Wallis test, n > 300 NMJs from N = 3 mice each for panel I. Data: mean ± SEM

Curious to ascertain if disease suppression is limited to the neuromuscular system or extends to other organs too, we also examined liver tissue in PND15 controls and SMA mutants with and without the modifier; hepatic pathology has been reported in SMA model mice as well as human patients and is characterized by fatty liver and pronounced microvesicular steatosis^20–23^. Expectedly, liver tissue of *Smn^2B/-^* mutants displayed disrupted architecture and widespread microvesicular vacuolization, consistent with severe steatosis. In contrast, liver sections from *Smn^2B/-^;Hspa8^G470R^* mice revealed a normal cellular architecture, uniform hepatocyte morphology, and complete absence of steatosis (Fig. S2). Collectively, these results demonstrate that Hspa8^G470R^ does not require the presence of the *SMN2* gene to suppress the SMA phenotype. Moreover, disease suppression in *Smn^2B/-^* mutants is not restricted just to the neuromuscular system but observed in peripheral tissue too.

### Hspa8^G470R^ effects SMN increase by retarding protein decay

Considering how robustly Hspa8^G70R^ protected against SMA, we sought to identify mechanisms of action of the modifier. *Smn^2B/-^* mice do not harbor *SMN2*. However, because the *Smn^2B^* allele was engineered to behave like *SMN2* and generate a transcript lacking murine *Smn* exon 7, and given the splice-switching property of Hspa8^G470R^, (refs. 14, 24) we commenced our assessment by examining *Smn^2B^* splice products in the various mouse cohorts. Predictably, healthy *Smn^2B/+^* controls produced copious quantities of FL (full-length) murine *Smn* – from both the wild-type (WT) and 2B alleles – and lower amounts of transcript lacking exon 7 – exclusively from the mutant allele (Figs. 3A, B). Moreover, as expected, the relative abundance of the two transcripts was reversed in *Smn^2B/-^* mutants. This ratio did not change in the presence of Hspa8^G470R^, (Figs. 3A, B and Fig. S3A) suggesting that notwithstanding having engineered the *Smn^2B^* allele to behave like *SMN2*, it is not subject to splice-switching by the modifier. We nevertheless proceeded to assess SMN protein levels in the different mouse cohorts. Consistent with past studies and the lower abundance of FL murine *Smn* transcript in *Smn^2B/-^* mutants, we found greatly reduced SMN protein in these mice relative to that in healthy controls (Figs. 3C, D). Protein levels in Hspa8^G470R^-expressing mutants remained very low but, surprisingly, did increase. The increase was modest or statistically insignificant in mutants heterozygous for the modifier but became noticeable – and significant – in mutants homozygous for Hspa8^G470R^ (Figs. 3C, D and Fig. S3B). Nevertheless, mean protein amounts in *Smn^2B/-^;Hspa8^G470R/G470R^* mutants never exceeded 25% of WT levels.

**Figure 3.**
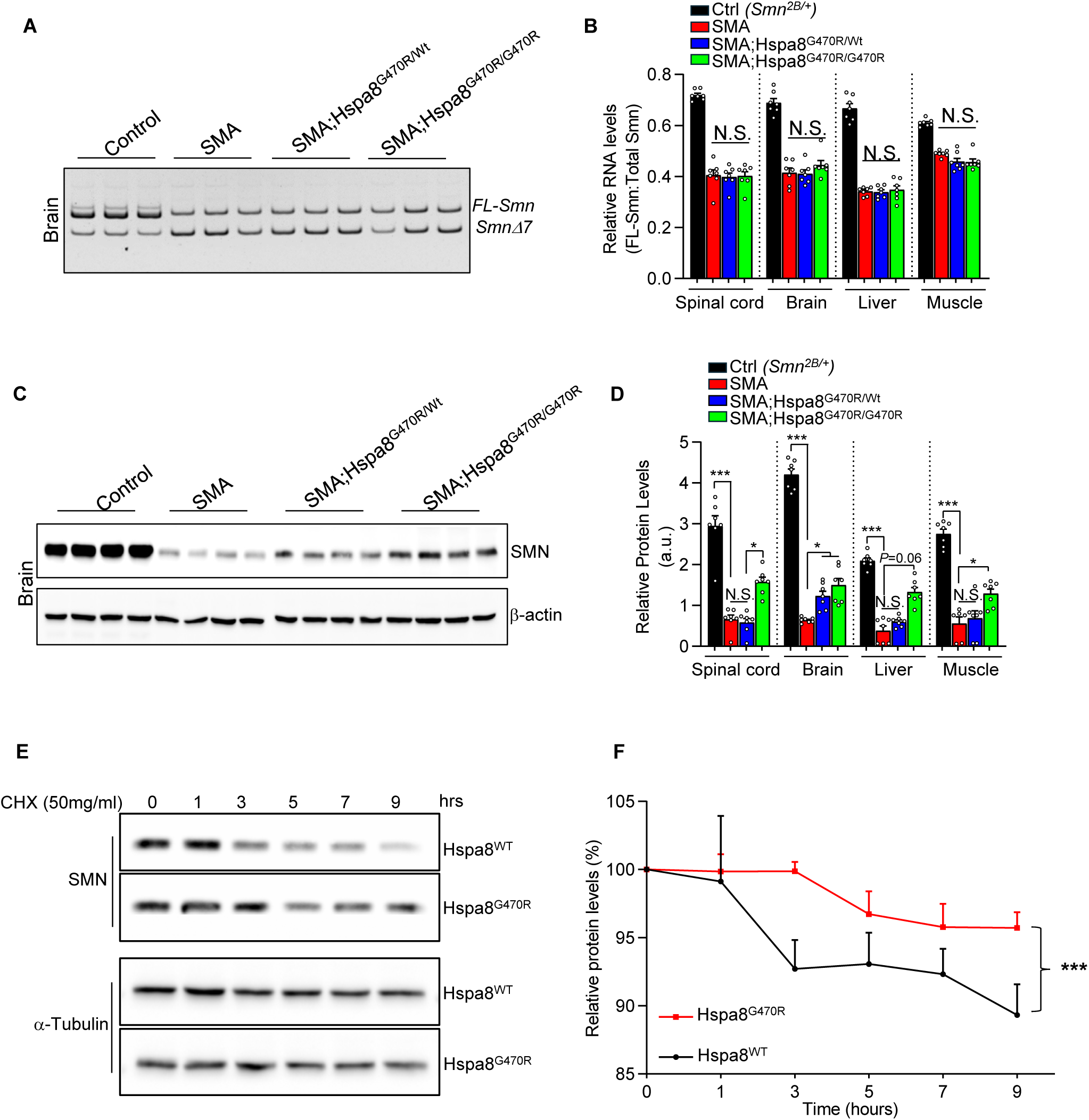
The Hspa8^G470R^ variant raises SMN levels by slowing protein decay. **(A)** Representative polyacrylamide gel electrophoresis of the FL and Δ7 RNA isoforms of the *Smn^2B^* allele in controls and SMA mutants with or without the Hspa8^G470R^ disease modifier. **(B)** Quantified results of the two RNA isoforms in diverse tissues of the four cohorts of mice. Kruskal-Wallis tests, N = 7 mice of each cohort. **(C)** Representative western blot of SMN protein in brain tissue of controls and mutants expressing either the WT or variant form of Hspa8. **(D)** Quantification of SMN protein in diverse tissues of the four cohorts of mice. *, ***, *P* < 0.05 and *P* < 0.001, respectively, one-way ANOVA (muscle and brain samples), Kruskal-Wallis test (liver and spinal cord samples), N = 7 mice of each genotype. **(E)** Representative western blot of SMN protein at indicated time point in MEFs expressing either Hspa8^WT^ or Hspa8^G470R^ following treatment with cycloheximide (CHX) to halt protein synthesis. **(F)** Graphical representation of SMN protein decay in MEFs either WT for Hspa8 or expressing the variant form of it. Note: ***, *P* < 0.001, AUC, results compiled from 3 independent experiments. Data: mean ± SEM

Considering unaltered FL murine *Smn* transcript levels but increased SMN protein in mutants harboring Hspa8^G470R^, we hypothesized a post-translational effect of the modifier. To test this, we assessed SMN turnover in mouse embryonic fibroblasts (MEFs) expressing either WT Hspa8 or the G470R variant. We found that SMN decay was indeed slowed in the presence of the variant (Figs. 3E, F). Moreover, consistent with a modifier-mediated effect on SMN *protein*, Hspa8^G470R^ did not affect murine *Smn* transcript stability (Fig. S3C). These results extend our prior findings demonstrating an interaction between SMN and Hspa8 (ref. 14) and reveal the physiological relevance of the interaction, which has been independently verified^25^.

### Hspa8 regulates SMN levels via chaperone-assisted selective autophagy (CASA)

Having established that Hspa8^G470R^ stabilizes the SMN protein likely via a direct physical interaction, we explored how such stabilization might be effected. To do so, we first investigated how the two proteins interact. This was accomplished by generating a series of deletion constructs expressing specific regions of the two proteins and using these in co-immunoprecipitation (co-IP) assays to identify minimal interacting domains on each. Expectedly, an intact, WT Hspa8 protein bound efficiently to bead-immobilized FL-SMN. Moreover, consistent with previous findings examining the interaction between the variant and human protein^14^, the affinity of the chaperone for murine SMN was reduced by the G470R variant (Figs. S4A – C). We also found that the N-terminal ATPase and nucleotide binding domain (NBD) of Hspa8 failed to bind SMN, whereas truncated products containing the chaperone’s substrate binding domain (SBD) did (Figs. S4A, B); deleting the C-terminal 104 amino acids of Hspa8, which constitutes the lid domain of the protein did not preclude it from binding SMN. In a similar manner, we determined that the Tudor, the Gemin-binding and the proline-rich domains of SMN are important for interacting with Hspa8 (Figs. S4D, E). The somewhat paradoxical observation that the N-terminal domain of SMN, which contains the first 151 amino acids of the protein and excludes the polyproline region binds Hspa8 while the SMN^ΔP^ fails to do so suggests that in the absence of the proline-rich region, the C-terminus of SMN sterically blocks Hspa8-SMN binding; neither region is present in the SMN_N construct, possibly enabling binding of the corresponding truncated protein to Hspa8. Notwithstanding this final observation, the overall binding data are consistent with SMN being one of several Hspa8 clients that interact with the latter’s SBD^25^.

Hspa8 is best known for ensuring cellular homeostasis through the sequestration or degradation of nascent, misfolded or aggregated proteins via autophagic pathways^26,27^. Coincidentally, there are several reports not just of perturbed autophagy in SMA, but that SMN itself is subject to autophagic degradation^28–33^. Prompted by these reports and our own observations above, we investigated autophagy as a potential link between SMN and Hspa8^G470R^. To do so, we first subjected WT MEFs to activators or inhibitors of autophagy and examined the effect of such treatment on SMN protein. Consistent with the notion that SMN is at least partly turned over by autophagy and that such turnover is mediated by the autophagy adaptor, p62, we found that serum starvation, an activator of autophagy, reduced the levels of both proteins, and SMN significantly so, whereas a leupeptin + NH_4_Cl (NL) cocktail, which inhibits lysosomal degradation, raised their concentrations (Figs. 4A - C). Interestingly, the decrease in SMN in serum-starved WT fibroblasts was mitigated in similarly treated cells expressing Hspa8^G470R^, a result congruent with an inhibitory effect of the variant on autophagic degradation of SMN. This resistance to SMN degradation was also observed in variant-expressing cells treated with the protein synthesis inhibitor, cycloheximide – relative to similarly treated WT cells. Predictably, SMN reduction in serum-starved WT cells treated with cycloheximide was greater than it was in the same cells cultured without drug. These experimental results bolster earlier reports of autophagic control of SMN^30–33^ and prompted us to investigate how Hspa8 might modulate this process.

**Figure 4.**
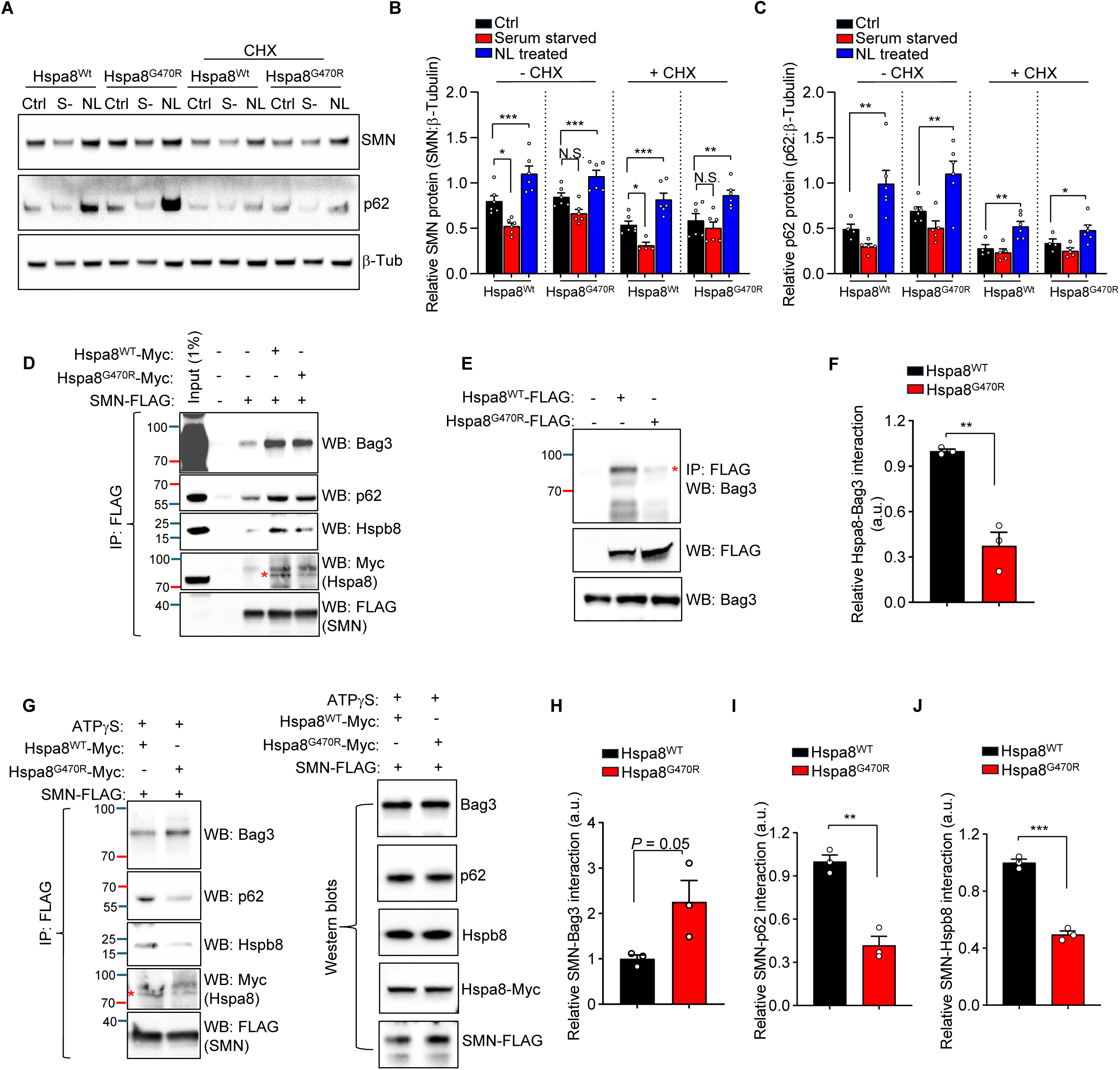
The Hspa8^G470R^ variant regulates SMN via chaperone-assisted selective autophagy. **(A)** Representative western blot of MEFs of indicated genotypes left untreated or treated to induce (serum starvation = S-) or inhibit (NL) autophagy. Quantification of **(B)** SMN and **(C)** p62 in fibroblasts depicted in panel A. Note: *, **, ***, *P* < 0.05, *P* < 0.01 and *P* < 0.001 respectively, one-way ANOVA, n = 6 replicates. **(D)** Representative co-IP and immunostains of CASA complex components bound to SMN in HEK293 cells co-transfected with either WT Hspa8 or the G470R variant of the chaperone. **(E)** Representative co-IP and western blot depicting reduced affinity of Bag3 for Hspa8^G470R^ compared to WT Hspa8. **(F)** Quantification of Bag3 protein bound to WT or the variant form of Hspa8. Note: **, *P* < 0.01, *t* test, n = 3 replicates. **(G)** Representative co-IP and immunostains of CASA component proteins pulled down with SMN in the presence of either the WT or the G470R variant form of Hspa8, following treatment with ATPγS. Right hand panel shows blots of endogenous or transfected proteins from cells used for the IPs. Quantified amounts of **(H)** Bag3, **(I)** p62 and **(J)** Hspb8 bound to SMN in the presence of WT or the G470R variant form of Hspa8 in experiments represented by panel G. Note: Comparisons made using the *t* test, n = 3 independent experiments. Also note: **, ***, *P* < 0.01 and *P* < 0.001 respectively; asterisks in panels D and G denote Hspa8-Myc bands and distinguish them from the larger, non-specific band in the gels. Data: mean ± SEM

Hspa8-mediated autophagy is effected several different ways^34^. We hypothesized that chaperone-assisted selective autophagy (CASA), a form of macroautophagy was the most likely pathway regulating SMN homeostasis, as the two other well-known forms of autophagy involving Hspa8, chaperone-mediated autophagy (CMA) and endosomal microautophagy (eMI), both require a KFERQ pentapeptide recognition motif on client proteins^34^; this motif is not found on SMN. Hspa8-assisted selective autophagy involves a large (CASA) complex, comprising among others the small heat shock protein, Hspb8, a nucleotide exchange factor (NEF) and co-chaperone such as Bag3 and the Hspa8 interactor and E3 ubiquitin ligase, Chip/Stub1 (ref. 27). The assembly of the complex and its interaction with client proteins is summarized in cartoon form (Fig. S5A). To test the idea that SMN proteostasis involves CASA, SMN-FLAG was immobilized on agarose beads and immunoprecipitates from cells co-transfected with either WT Hspa8-Myc or a tagged version of the chaperone variant examined for evidence of CASA complex members. Predictably, Hspa8 was pulled down with SMN, and the G470R variant found to bind less efficiently than WT Hspa8 to SMN. However, we also found copious levels of endogenous Bag3 in the immunoprecipitates and detectable but lower amounts of endogenous Hspb8 and p62 (Fig. 4D). This indicates that SMN does indeed interact with CASA complex components. Because binding of Bag3 to Hspa8 regulates ADP to ATP exchange on the chaperone and thus determines how efficiently client proteins like SMN are released by the chaperone as they progress through the autophagic pathway (Fig. S5), we also sought to examine if and how Hspa8^G40R^ affects Hspa8-Bag3 interaction. Accordingly, we immunoprecipitated tagged WT Hspa8 or Hspa8^G470R^ and quantified levels of endogenous Bag3 that were pulled down by the respective proteins. Expectedly, Bag3 was pulled down with WT Hspa8. Interestingly, however, it bound significantly less well to Hspa8^G470R^ (Figs. 4E, F). This suggests that client proteins like SMN, once bound to Hspa8^G470R^, are unlikely to be efficiently released as refolded proteins or, in instances where refolding fails, to phagosomes for degradation, instead becoming entrapped within the closed-conformation of the chaperone (Fig. S5B). To test this idea and furthermore avoid variability in interactions resulting from dynamic changes between ADP/ATP-bound Hspa8 states that can be affected by small changes in buffer or temperature, we repeated the SMN-FLAG immunoprecipitations with co-transfected Hspa8^G470R^ or WT Hspa8. However, we did so following treatment of cells with ATPγS, a modified form of ATP that is slow to hydrolyze, locks Hspa8 into an ATPγS-bound state and is therefore suitable for detecting intermediate CASA complexes. Under this condition, not only did we detect significantly higher amounts of the SMN-Hspa8-Bag3 complex when cells over-expressed the G470R variant but also reduced amounts of SMN-Hspb8 and SMN-p62 complexes in the presence of the variant (Figs. 4G – J). A rise in the SMN-Hspa8^G470R^-Bag3 complex suggests that SMN does indeed become entrapped within this CASA intermediate. Congruently, reduced binding of SMN to Hspb8 and the autophagosome adaptor p62 in the presence of the G470R variant suggests that SMN is inefficiently released from the SMN-Hspa8^G470R^-Bag3 complex for lysosomal degradation. The net outcome is a modest but disease-relevant increase in SMN. Overall, our results shed additional light on SMN regulation via a specialized form of macroautophagy that involves the CASA complex. In so doing, they assign a second, novel SMA-modifying mechanism of action to the Hspa8^G470R^ chaperone variant.

### The Hspa8^G470R^ variant enhances synaptic strength and the efficiency of synchronous NMJ neurotransmission

Considering the improved agility of *Smn^2B/-^;Hspa8^G470R^* mutants and our prior study^14^ demonstrating that the variant contributes to motor recovery at least in part by stimulating neuronal SNARE complex formation and normalizing neurotransmission, we inquired if and how well Hspa8^G470R^ had mitigated synaptic dysfunction in *Smn^2B/-^* mice. To do so, we focused on PND15 NMJs of the transverse abdominus (TVA), a muscle relevant to the righting reflex we employed (Fig. 1C) to assess motor function. We found that neither the amplitudes nor frequencies of miniature endplate potentials (mEPPs) in *Smn^2B/-^* mutants were altered compared to those of *Smn^2B/+^* controls; the presence of Hspa8^G470R^ in the mutants did not change these values, although mEPP frequency was modestly increased in controls heterozygous for the modifier (Figs. S6A, B). Expectedly, however, evoked potential (EPPs) and quantal content *(m)* in *Smn^2B/-^*mutants were significantly lower (65% and 54% respectively) than those of controls (Figs. 5A, B). Notably, Hspa8^G470R^ raised these measures in mutants, with the increases not only became highly significant in *Smn^2B/-^;Hspa8^G470R/G470R^* mice – relative to *Smn^2B/-^* mice – but also, surprisingly, surpassing values in healthy *Smn^2B/+^* controls (Figs. 5A, B and Figs. S6C, D). Additionally, Hspa8^G470R^ also raised EPPs and quantal content in healthy controls (Figs. S6E, F). Together, these results suggest that Hspa8^G470R^ is a potent effector of NMJ neurotransmission. Moreover, considering its ability to potentiate neurotransmission even in healthy controls and its propensity to raise EPPs and quantal content in *Smn^2B/-^;Hspa8^G470R/G470R^* mutants beyond those of *Smn^2B/+^* mice – despite the markedly lower SMN levels in the former cohort (Figs. 3C, D) – our results suggest that the modifier’s effects on NMJs are direct and not merely because it stabilizes and raises SMN.

**Figure 5.**
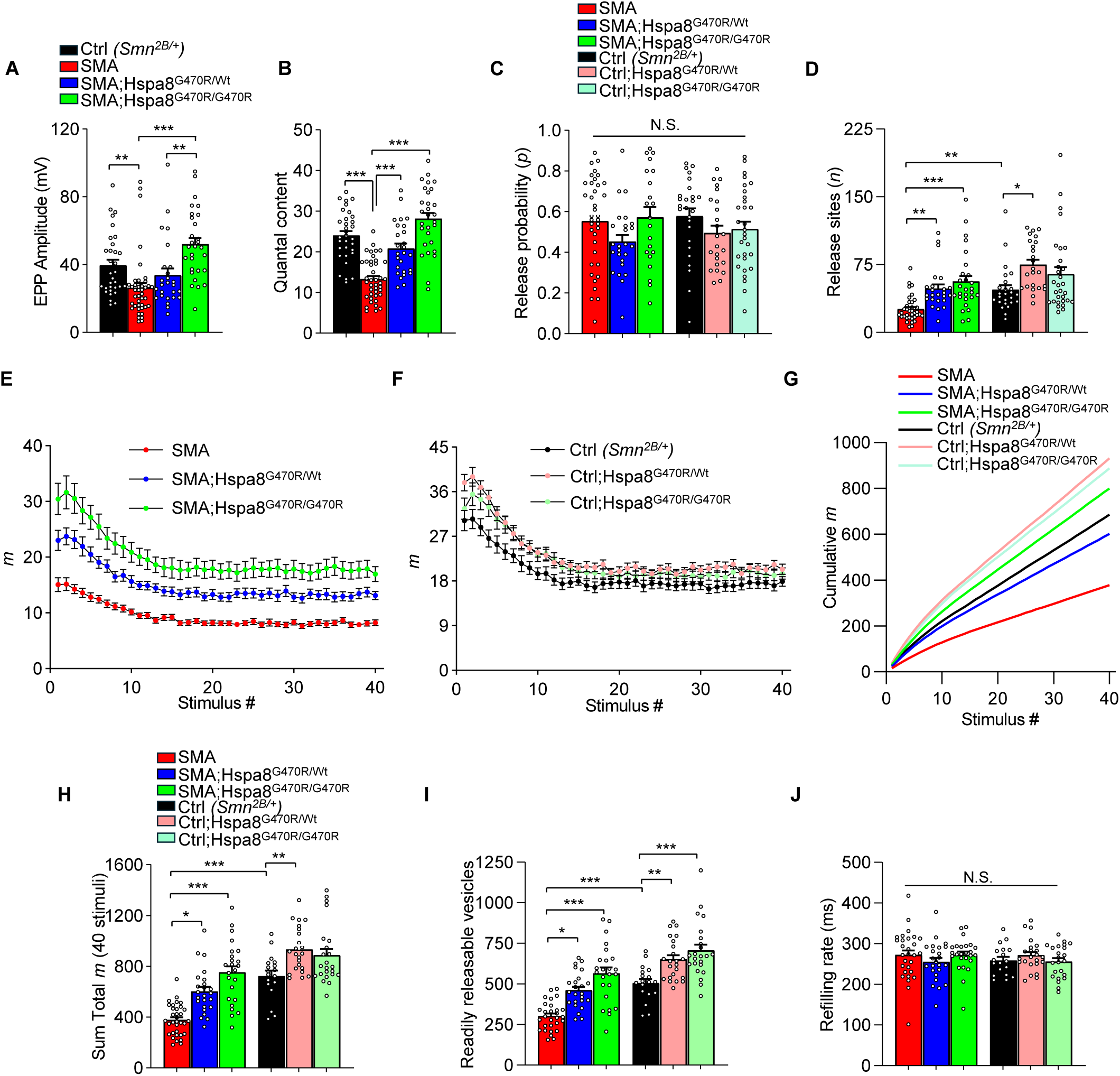
Hspa8^G470R^ potentiates neurotransmission in SMA and control mice. Quantified results of **(A)** evoked potentials and **(B)** quantal content in PND15 controls and SMA mice with or without the G470R variant. Note: **, ***, *P* < 0.01 and *P* < 0.001 respectively, Kruskal-Wallis test (panel A) and one-way ANOVA (panel B). Estimates of **(C)** release probability and **(D)** number of neurotransmitter release sites at NMJs of SMA and control mice with or without Hspa8^G470R^. Note: *, **, ***, *P* < 0.05, *P* < 0.01 and *P* < 0.001 respectively, one-way ANOVA (panel C) and Kruskal-Wallis test (panel D). Mean quantal content at each of 40 stimuli in response to high-frequency stimulation in **(E)** SMA mutants and **(F)** controls with 0, 1 or 2 copies of the Hspa8^G470R^ modifier. Note: means estimated from n ≥ 25 NMJs from N = 3 – 4 mice of each cohort. **(G)** Traces depicting accumulating quantal content over 2.5s of the high-frequency stimulation in each of the six cohorts of mice analyzed in panels E and F. Note higher quantal content in mutants and controls with the modifier relative to their respective cohorts expressing Hspa8^WT^. **(H)** Cumulative quantal content in the various mouse cohorts at the conclusion of the train. Note: *, **, ***, *P* < 0.05, *P* < 0.01 and *P* < 0.001 respectively, Kruskal-Wallis test. **(I)** Hspa8^G470R^ increases the RRP of NMJ vesicles in SMA and control mice. Note: *, **, ***, *P* < 0.05, *P* < 0.01 and *P* < 0.001 respectively, Kruskal-Wallis test. **(J)** Neurotransmitter refilling rates in the various mouse cohorts remain unaltered by the modifier; Kruskal-Wallis test. Note: All quantifications here are based on n ≥ 25 NMJs from N = 3 – 4 mice of each cohort. Data: mean ± SEM

The potentiated neurotransmission we observed at NMJs expressing the Hspa8^G470R^ variant could derive from increased vesicle release probability *(p)*, a greater number of functional release sites *(n)*, or both. To estimate which, if either, mechanism might be responsible for the enhanced synaptic function in the presence of the modifier, we employed a simplified binomial model of synaptic transmission in which *m* = *p* * *n* (Ref. 35). This assessment failed to reveal differences in release probability in the various mouse cohorts whether or not they expressed Hspa8^G470R^ (Fig. 5C). In contrast, we found that *n* values at mutant NMJs rose significantly in the presence of the modifier and, in fact, were normalized to control *(Smn^2B/+^)* levels (Fig. 5D). This effect of Hspa8^G470R^ on *n* was observed at control *(Smn^2B/+^)* NMJs too, raising the number of functional release sites at these synapses to supraphysiological levels and mirroring outcomes of EPP and quantal content estimates (Fig. 5D). To show experimentally that the calculated increase in *n* by Hspa8^G470R^ did indeed reflect greater numbers of effective neurotransmitter release sites, we subjected the NMJs of the various cohorts of mice to high-frequency (20 Hz, 2.5s) nerve stimulation trains. Under this protocol, EPP values experience a gradual decrement and then stabilize, reflecting the dynamics – vesicle mobilization, depletion and replenishment – of the RRP. Akin to outcomes following low-frequency (0.5Hz) stimulation (see Figs. 5C, D), we found that neurotransmitter release in response to high-frequency trains was consistently higher at SMA NMJs in the presence of Hspa8^G470R^ (Fig. 5E). This pattern of enhanced release was also seen at control NMJs (Fig. 5F). Plots of cumulative neurotransmitter release (Fig. 5G), the sum total *m* (Fig. 5H) and plateau amplitudes (Fig. S6G) reflected these findings, confirming the estimated *n* values (see Fig. 5D) and suggesting that even a single Hspa8^G470R^ allele is sufficient to restore neurotransmission in *Smn^2B/-^* mutants to *Smn^2B/+^* control levels.

What is the physiological correlate of increased *n* in the presence of Hspa8^G470R^ and how might one explain the variant’s effects during high-frequency stimulation? We first investigated possible alterations in short-term plasticity. However, we did not observe significant differences in either paired-pulse facilitation (PPF) or short-term depression (STD) among *Smn^2B/-^* or *Smn^2B/+^* mice expressing zero, one, or two copies of the modifier (Figs. S6H, I). An alternative explanation for increased neurotransmission in the presence of the G470R variant is an increase in the effective size of the RRP of vesicles. The RRP size is critically influenced by the availability and assembly of SNARE complexes, which mediate vesicle docking and confer fusion competence^36^. Importantly, Hspa8 is an established component of a tripartite chaperone complex known to be important for SNARE complex assembly, and the G470R variant is known to stimulate the formation of such SNARE complexes^14,16^. Accordingly, we investigated functional RRP size in the presence or absence of Hspa8^G470R^. To do so while simultaneously assessing the dynamics of vesicle mobilization, depletion and replenishment, we employed a sequential kinetic model^37,38^. This model (Figs. S7A, B) posits that docked vesicles in the RRP are the first to be released during repetitive stimulation, that the pool undergoes exponential depletion, and that vesicle recruitment increases over time to sustain release^39^. Predictably, we estimated that RRP size was substantially lower in *Smn^2B/-^*mutants than it was in healthy controls. Significantly, Hspa8^G470R^ restored this to control levels, with RRP size rising in a dose-dependent manner (Fig. 5I). Equally notably, the variant similarly raised RRP size in controls (Fig. 5I), emphasizing its broad impact on synaptic vesicle availability. An increase in effective RRP size by Hspa8^G470R^ might be effected by inducing SNARE complex assembly or slowing disassembly of these complexes, e.g., when N-ethylmaleimide-sensitive factor (NSF) is inhibited^36^. To rule out impairments in SNARE complex *disassembly*, we estimated the mean vesicle refilling rate in *Smn^2B/-^* and *Smn^2B/+^* mice with or without the modifier. We found that refilling rates remained unchanged irrespective of genotype and modifier copy number (Fig. 5J). These results imply that the modifier does not significantly affect the kinetics of SNARE complex disassembly, endocytosis, or the recruitment of new vesicles to release sites. Instead, the observed RRP increase is most likely due to enhanced SNARE complex assembly by the G470R variant. Collectively, our physiological assessments of neuromuscular activity bolster notions of a direct, SMN-independent effect of Hspa8^G470R^ on synaptic strength and NMJ function.

### Hspa8^G470R^ from an exogenous source mitigates the SMA phenotype in model mice

Rescue of the SMA phenotype when Hspa8^G470R^ is genetically expressed raises the possibility of using this specific variant or targeting endogenous Hspa8 to treat the disease. To test this idea, we packaged an expression cassette consisting of the modifier under the control of a chicken β-actin promoter into AAV-PHP.eB, a serotype that transduces neurons especially robustly^40^, and introduced it, or vehicle, via the cerebral ventricles into PND0 *Smn^2B/-^* mutants. Predictably, vehicle-treated mutants began exhibiting overt disease at two weeks of age and succumbed to SMA by the third week of life. In contrast, median survival of AAV-PHP.eB-Hspa8^G470R^-treated mutants increased by ∼100% to 33 days; the longest living mutants survived beyond 100 days (Fig. 6A). Consistent with this observation, overall health of the mutants receiving AAV-PHP.eB-Hspa8^G470R^, as assessed by body weight, was significantly improved by the second postnatal week of life (Fig. 6B and Figs. S8A, B) relative to vehicle-treated *Smn^2B/-^* cohorts. This was further reflected in greater agility of these mutants (Fig. 6C). Additionally, we found that overt phenotypic rescue following delivery of AAV-PHP.eB-Hspa8^G470R^ to the mutants was accompanied by significantly less pathology at nerve-muscle connections. At PND18, mean endplate size in mutants receiving the Hspa8^G470R^ construct was significantly larger than that of age-matched vehicle-treated mutants in both the proximal triceps as well as the distal gastrocnemius muscles (Fig. 6D). Correspondingly, denervation of the muscles was suppressed (Figs. 6E, G) and the characteristic, abnormal accumulation of NF protein in SMA nerve terminals markedly reduced (Figs. 6F, G). While the overall rescue of disease in AAV-PHP.eB-Hspa8^G470R^-treated mutants is more modest than that seen when the variant is expressed endogenously, these results nevertheless constitute compelling evidence that the modifier has a potent SMA-suppressing effect even when expressed from an exogenous source delivered postnatally.

**Figure 6.**
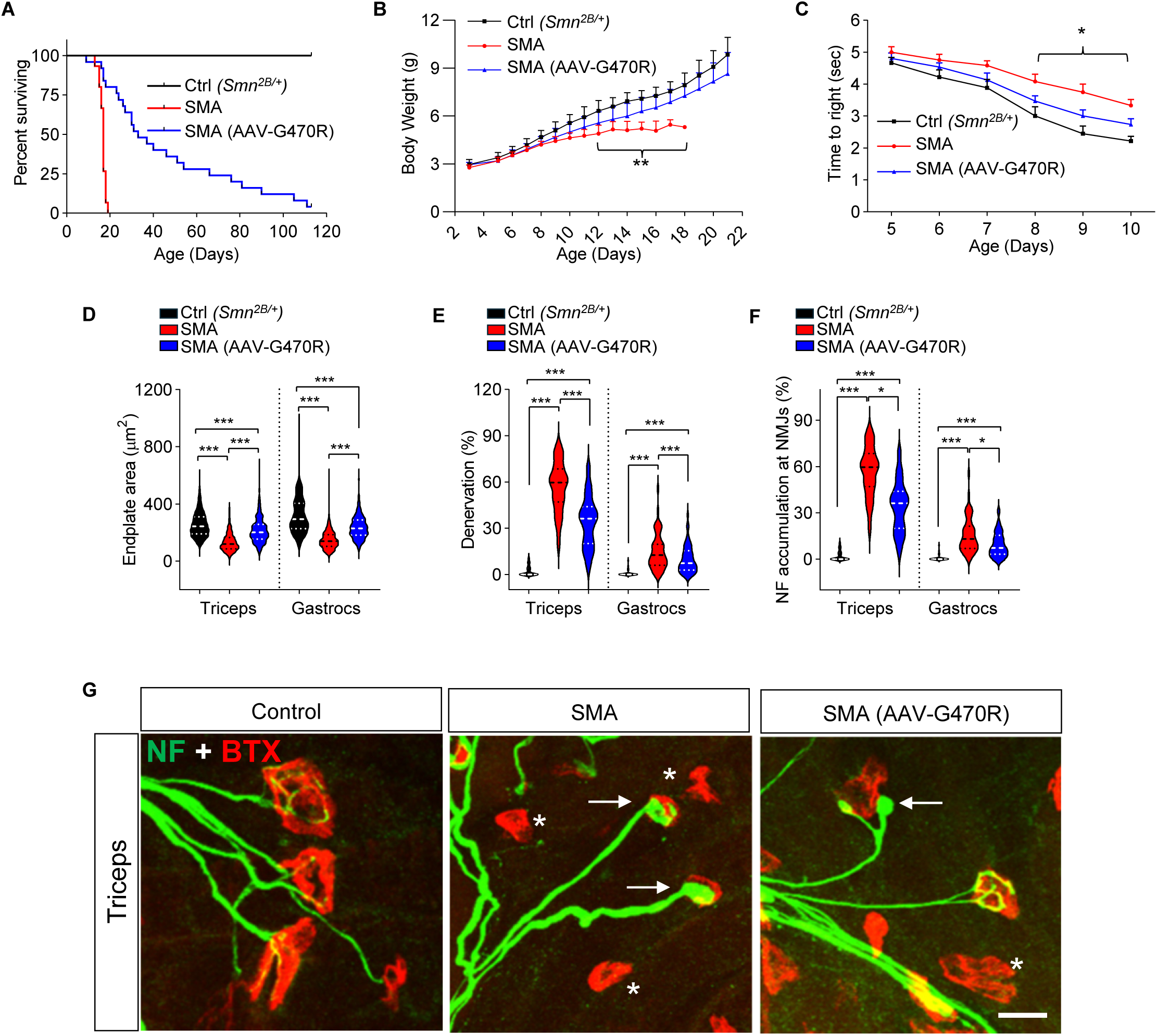
AAV-mediated delivery of Hspa8^G470R^ mitigates disease in SMA mice. (A) Kaplan-Meier survival curves depicting increased lifespan of SMA mutants administered AAV-PHP.eB-Hspa8^G470R^. *P* < 0.0001 between vehicle-treated and AAV-treated mutants, log-rank test, n = 10 – 25 mice of each cohort. Mutants treated with AAV-PHP.eB-Hspa8^G470R^ (B) gain significantly more weight and (C) perform better in the righting reflex assay than vehicle-treated SMA mice. Note: *, **, *P* < 0.05, *P* < 0.01, *t* tests between the two groups, n = 10 – 23 mice. Quantified (D) mean NMJ size, (E) degree of denervation and (F) pathology in nerve terminals assessed by NF swellings in the three cohorts of mice. Note: *, **, ***, *P* < 0.05, *P* < 0.01 and *P* < 0.001 respectively, Kruskal-Wallis tests (panels D, F) and one-way ANOVA (panel E), n = 500 – 800 AChR clusters from N = 4 mice of each cohort. (G) Representative photomicrographs of NMJs in triceps muscle from healthy controls or mutants administered either AAV-PHP.eB-Hspa8^G470R^ or vehicle. Asterisks denote denervated endplates; arrows highlight relatively simplified nerve terminals abnormally swollen with NF protein. Note fewer such terminals and denervated endplates in muscle from the mutant administered AAV-Hspa8^G470R^. Scale bar – 20μm. Data: mean ± SEM

### Delayed-onset neuromuscular disease, in Smn^2B/-^;Hspa8^G470R/Wt^ mutants, models intermediate (type II) SMA

One copy of the Hspa8^G470R^ allele mitigates the severe phenotype of SMNΔ7 SMA model mice but fails to prevent disease onset before weaning or extend lifespan beyond 4 weeks of age^14^. In contrast, we were unable to detect overt disease in 2-week-old *Smn^2B/-^;Hspa8^G470R/Wt^* mice (Fig. 1B, C). Still, because our anatomical studies uncovered neuromuscular pathology in these mutants, we continued to monitor them closely through early adulthood. At ∼4 months of age, we did not detect any difference in forelimb strength in the mutants (Fig. 7A). Yet, ambulation was perturbed in a proportion of these animals and visually linked to altered hindlimb function. This defect worsened until paralysis of the hindquarters occurred between 5 and 6 months of age (Supplemental Video 1). Altered gait by 4 months of age in these mutants was reflected in an abnormal hindlimb splay reflex, whereas no such defect was detected in mutants homozygous for Hspa8^G470R^ (Fig. 7B); ambulation in Hspa8^G470R^ *homozygous* mutants remained normal even when their heterozygous counterparts were paralyzed (Supplemental Video 1).

**Figure 7.**
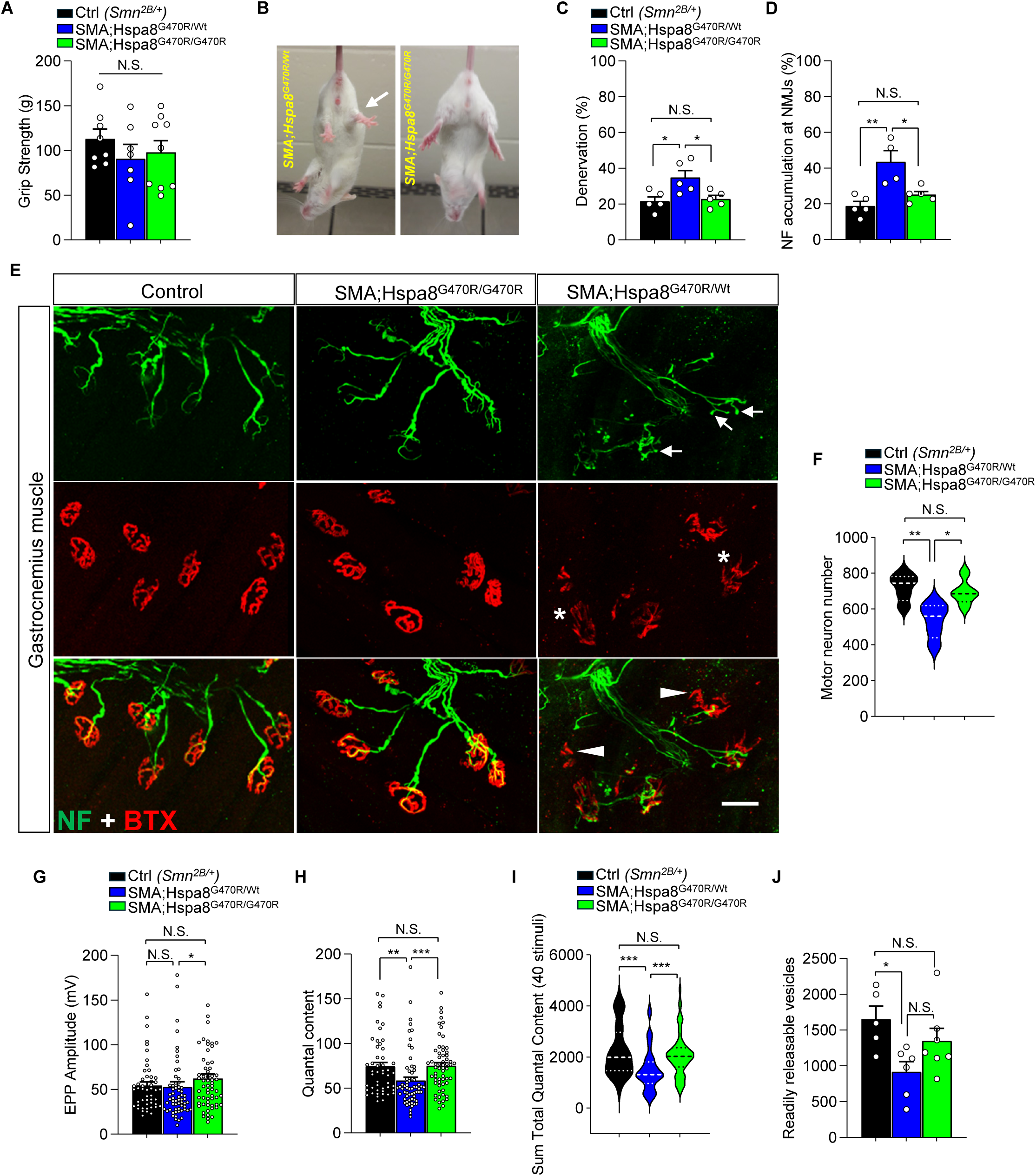
Delayed-onset disease in *Smn^2B/-;^Hspa8^G470R/Wt^* mutants models type II SMA. **(A)** Forelimb function remains unaffected in 4-month-old mutants bearing the G470R variant. Comparisons done using one-way ANOVA, n = 7 – 9 mice. **(B)** A reduced hindlimb splay reflex (arrow) in 4-month-old mutants with one but not two copies of the variant signifies neuromuscular weakness in these limbs. Four-month-old mutants heterozygous for Hspa8^G470R^ exhibit greater **(C)** denervation and **(D)** nerve terminal pathology. Note: *, **, *P* < 0.05 and *P* < 0.01 respectively, one-way ANOVA, n = 135 NMJs from N = 5 mice of each cohort. **(E)** Representative NMJs in 4-month-old controls and mutants bearing one or two copies of Hspa8^G470R^. Note denervation (arrowheads), fragmented endplates (asterisks) and simplified nerve terminals in the form of endbulbs (arrows) in the mutant with one G470R variant allele. Scale bar – 20μm. **(F)** Graph of morphometric counts of lumbar spinal motor neurons in 4-month-old controls and mutants with the modifier. Note: *, **, *P* < 0.05 and *P* < 0.01 respectively, one-way ANOVA, N = 5 mice of each cohort. Quantified results of NMJ electrophysiology performed at 0.5Hz stimulation on the three mouse cohorts depict **(G)** unaltered EPPs and **(H)** reduced quantal content in mutants heterozygous for the modifier. Note: *, **, *P* < 0.05, *P* < 0.01, Kruskal-Wallis tests (panels G, H). **(I)** Sum total of quantal content values at the end of 40 stimuli along a high-frequency (20Hz, 2.5s) stimulation train depicts a significantly lower value in 4-month-old mutants heterozygous for the G470R variant relative to the two other mouse cohorts. Note: ***, *P* < 0.001, Kruskal-Wallis test. **(J)** Fewer readily releasable vesicles were estimated in SMA mice heterozygous for the modifier. Note: *, *P* < 0.05, one-way ANOVA. Also note that samples sizes of n = 50 – 60 NMJs from N = 5 – 6 mice were employed (panels G – J). Data: mean ± SEM.

Hindlimb weakness and paralysis are frequently preceded by neuromuscular pathology. Expectedly, an examination of NMJs in the gastrocnemius of *Smn^2B/-^;Hspa8^G470R/Wt^* mutants at 4 months revealed significantly greater denervation accompanied by abnormal amounts of NF protein in nerve terminals (Figs. 7C – E). Interestingly, AChR clusters in these mutants were also frequently disassembled and fragmented (Fig. 7E), no longer conforming to the normal, elaborate pretzel-shaped endplates we observed in healthy controls and mutants homozygous for the modifier. Consistent with the NMJ defects, we found fewer motor neurons in the lumbar spinal cord of 4-month-old mutants with a single Hspa8^G470R^ allele (Fig. 7F); mutants with two copies of the modifier retained a normal complement of spinal motor neurons. Electrophysiological analysis of NMJ function in the TVA muscle of these two cohorts of mice and age-matched controls reflected many of our anatomical findings. At low frequency (0.5Hz)stimulation, mean evoked potentials trended lower in *Smn^2B/-^;Hspa8^G470R/Wt^* mice but were not statistically different from those in controls (Fig. 7G). In contrast, quantal content in these mutants did differ significantly from control values and was markedly lower (Fig. 7H). mEPP amplitudes were not altered in mutant mice, but interestingly, mEPP frequency was (Figs. S9A, B). The lower quantal content assessed at 0.5Hz stimulation was also apparent in response to high-frequency (20 Hz, 2.5 s) trains (Fig. 7I and Figs. S9C, D). Finally, we inquired if and how Hspa8^G470R^ had affected short-term plasticity and the RRP in young adult mutants. We found that whereas RRP in mutants homozygous for the G470R variant remained normal, RRP in mutants heterozygous for the modifier was statistically lower than control values (Fig. 7J). Paired-pulse facilitation remained unaltered in the mutants (Fig. S9E). However, interestingly, short-term depression rose in both sets of animals relative to that estimated in healthy controls (Fig. S9F), implying altered neurotransmission. Collectively, these results suggest that notwithstanding a relatively normal phenotype assessed at two weeks of age, mutants with just one copy of the modifier develop late-onset neuromuscular disease accompanied by functional and anatomical deficits of the NMJs. The steady deterioration of neuromuscular health in these mutants between PND14 and ∼6 months of age, followed by a slower decline that resulted in several such animals surviving beyond 10 months is reminiscent of type II SMA, making this line of mice a unique resource to investigate intermediate forms of the human disease.

## Discussion

A clear understanding of the precise mechanisms linking low SMN protein to neuromuscular dysfunction in SMA has lagged the development of therapies for the disease. While the therapies, which focus primarily on SMN repletion, are major breakthroughs, having profoundly altered the prognosis for most newborns diagnosed with the disease, they have done comparatively little for patients who received treatment late. Moreover, in very severely affected patients, newborn treatment prevents early mortality but is increasingly associated with novel disease phenotypes^4–6^. These observations reinforce arguments for a renewed focus on basic SMA mechanisms. One means of revealing such mechanisms is through the identification of novel disease modifiers. Here, we report on one such modifier – a G470R variant of the widely recognized proteostasis factor and synaptic chaperone, Hspa8. Our overall results reveal new mechanisms of action of the modifier. Correspondingly, they shed additional light on the basic biology of SMN and specific pathways relevant to neuromuscular dysfunction in SMA. Five principal findings emerge from our study. First, we demonstrate that the modifier mitigates the SMA phenotype just as potently in model mice devoid of *SMN2* as it does in mutants bearing the *SMN1* copy gene. This attests to the *bona fide* SMA-modifying effects of Hspa8^G470R^ and demonstrates that its disease suppressing actions are possible without necessarily altering *SMN2* splicing. Second, we show that despite the absence of *SMN2*, disease modification is effected, at least in part, by augmenting the SMN protein. Our findings link the increase in the protein to effects of Hspa8^G470R^ on SMN turnover, specifically via autophagy-associated pathways. Third, our study provides additional evidence of a direct and SMN-independent effect of the SMA modifier in mitigating disease. This effect, likely related to the role of Hspa8 in chaperoning synaptic SNARE complexes, involves an increase in the readily releasable pool of synaptic vesicles and a corresponding potentiation of neurotransmission not only at SMA NMJs but also at healthy synapses. Enhanced neurotransmission in the presence of Hspa8^G470R^ or agents that similarly modulate Hspa8 function is potentially useful for myriad neuromuscular diseases and thus relevant beyond just SMA. Fourth, we provide proof-of-concept data of the feasibility of employing either Hspa8^G470R^ or factors that target Hspa8 as therapeutic agents. In our study, AAV-mediated expression of Hspa8^G470R^ robustly suppressed disease in model mice. Finally, and serendipitously, we have discovered that a single, G470R variant allele on the *Smn^2B/-^* background is sufficient to delay but not prevent neuromuscular dysfunction. Onset of disease in young adult mutants culminated in total hindlimb dysfunction by 6 months of age and constitutes, to our knowledge, the first truly paralytic model of SMA and thus a useful representation of intermediate disease in humans.

The *SMN2* splice-modulating function we previously attributed to Hspa8^G470R^ confounded inferring any potential direct and SMN-independent SMA-ameliorating effects of the modifier^14,41^. Accordingly, we sought to reveal such mechanisms by testing the modifier in *Smn^2B/-^* model mice, which are devoid of *SMN2*. Reassuringly, we found that the modifier also potently suppresses disease in such mice. Yet, unexpectedly, SMN protein once again rose modestly – not because of a gene splice-switching event but rather owing to reduced SMN turnover. Our results implicate chaperone-assisted selective autophagy (CASA) in this outcome and suggest that normal processing of SMN through this pathway is thwarted not only by weakened interactions between SMN and the G470R variant but also by altered affinity of the variant for the NEF and CASA co-chaperone, Bag3. Because this latter interaction is critically important for release of Hspa8 client proteins, either as a functional, refolded molecule or, in the instance that refolding fails, via the multi-protein CASA complex to the autophagy adaptor, p62, for lysosomal degradation^27,32^, we infer that SMN processing through the CASA pathway is retarded, and residual protein trapped in Hspa8^G470R^-SMN-Bag3 intermediate complexes. The apparently paradoxical relative increase in this complex vis-à-vis complexes containing WT Hspa8, considering reduced affinity of Hspa8^G470R^ for both SMN and Bag3 could be an effect of the variant on the overall 3-D structure of Hspa8, rendering its lid domain unable to respond appropriately to Bag3 binding, revert to an open conformation and release bound client protein. While additional work will be needed to reveal such details, it is clear from our work that SMN engages less with both Hspb8, a member of the CASA complex that is important for delivery of misfolded proteins to phagosomes, and p62 – observations strongly suggestive of reduced autophagy turnover of the protein. Collectively, these results have two important implications. First, they link Hspa8 to a second, SMN-modulating function – one directed at the *protein*. To our knowledge, this is the first factor to have dual effects on SMN, functioning not only as an *SMN2* splice-switcher but also serving to stabilize SMN protein. Second, they suggest that selectively targeting autophagy could be of important therapeutic value in the treatment of SMA.

The unexpected discovery that Hspa8^G470R^ can bypass *SMN2* and nevertheless raise SMN, once again confounds unequivocally assigning to the variant an SMN-independent effect in SMA suppression. Yet, our findings here strengthen notions of such an effect. Given its SMN-stabilizing role, rescue of the SMA phenotype of *Smn^2B/-^* mice is not surprising. However, the degree of phenotypic rescue – greater than10-fold increase in lifespan, particularly in *Smn^2B/-^;Hspa8^G470R/Wt^* mice – is. In these Hspa8^G470R^ heterozygous mutants, SMN generally did not exceed that expressed by *Smn^2B/-^* mutants, yet disease rescue was significant. In contrast, in a previous study which involved p62 knockdown in SMA mutants and resulted in an increase in SMN of ∼75%, lifespan of the mutants was merely enhanced by half^32^. Direct, Hspa8^G470R^-mediated effects at NMJs provide one explanation for the more potent disease-mitigating effects of the chaperone variant. Indeed, not only did the modifier potentiate neurotransmission significantly in SMA mutants but also did so in healthy controls. Moreover, direct comparisons of NMJ function in *Smn^2B/-^*;*Hspa8^G470R/G470R^* mutants and healthy *(Smn^2B/+^)* controls revealed a modest yet significant potentiation of neurotransmission in the former – notwithstanding markedly lower SMN levels in the mutants. Our results suggest that the enhanced neurotransmission derives less from modest increase of SMN by the modifier than from a primary, synapse-specific Hspa8^G470R^ role – in augmenting the readily releasable pool of synaptic vesicles. This pool of synaptic vesicles is known to be influenced by SNARE complex levels^36^, which in turn are critically dependent on an Hspa8-containing synaptic chaperone complex^15^. Modulation of the SMA phenotype via direct effects of Hspa8^G470R^ at NMJs on SNAREs or related factors whose activities are subject to fine-tuning by CASA would assign it a surprising third mechanism of action in mitigating the SMA phenotype. More generally, the effects of the variant on the NMJ would be relevant for motor neuron diseases at large.

What is the therapeutic potential of Hspa8^G470R^? Our AAV-mediated results clearly attest to its therapeutic potential. The more modest rescue we observed when the variant was virally delivered is a likely consequence of at least three factors. First, there is a well-established lag between administration of AAV constructs and their physiological effects^42^. In severe SMA mice, this mutes phenotypic rescue. Second, disease modification is influenced by AAV biodistribution; the level of vector spread and, consequently, payload expression is invariably inferior to the uniform expression from endogenous alleles. This too is expected to reduce treatment outcomes. Finally, we posit that the presence of endogenous WT Hspa8 in our mutants contributes to the sub-optimal disease-modifying effects of the variant by competing with the latter in relevant SMA-suppressing pathways. Future work will assess if this last handicap can be overcome and treatment outcomes improved by editing the endogenous *Hspa8* gene to bring about the single g1489c nucleotide conversion that underlies the G470R variant.

Considering the abolition of neuromuscular disease by Hspa8^G470R^ in the Δ7 line of SMA model mice^14^, an unexpected outcome of this study was the emergence of a late-onset phenotype, culminating in total hindlimb paralysis of *Smn^2B/-^*;*Hspa8^G470R/Wt^* mutants. Our analysis of these mutants indicated normal neuromuscular function at PND14. Yet, we did detect modest NMJ pathology in limb muscles of such mutants. These defects grew inexorably, particularly in hindlimbs, eventually causing complete hindlimb immobility by 6 months; age-matched mutants homozygous for the G470R variant remained asymptomatic and overtly normal, although we lost a small (∼15%) proportion of such animals at 12 months of age to undetermined causes. What accounts for the severe, albeit delayed-onset phenotype of *Smn^2B/-^*;*Hspa8^G470R/Wt^* mutants considering abolition of neuromuscular disease by the variant in Δ7 SMA mice? One intriguing explanation stems from the SMN alleles in the two mouse models. Δ7 SMA mice^17^ are endowed with 2 WT copies of human *SMN2* and therefore express low but fully functional WT FL-SMN protein. In contrast, *Smn^2B/-^* mutants do not express any WT SMN; the *Smn^2B^* allele is engineered to mis-splice but does so by relying on an Smn^G282F^ mutation in an exon splice enhancer of murine *Smn* exon 7 (ref. 43). In humans, this residue is conserved and corresponds to position 287 of the SMN protein. Although missense mutations at SMN^G287^ have not been linked unequivocally to SMA, the following residue at position 288 has (www.ncbi.nlm.nih.gov/clinvar/). Notably, G287 also lies within 7 amino acids of the functionally important YG box, which is implicated in SMN oligomerization^44–47^. Consequently, it is plausible that Smn^G282F^ in *Smn^2B/-^* mutants is a mild, oligomerization-incompetent protein whose effects at low concentrations only become apparent over time. The delayed onset phenotype of *Smn^2B/-^;Hspa8^G470R/Wt^* mice and the much milder phenotype of *Smn^2B/-^;SMN2^Tg/+^* mutants^48^ wherein WT FL-SMN is expected to oligomerize with and stabilize Smn^2B^ protein are consistent with this hypothesis. Notwithstanding these observations, *Smn^2B/-^;Hspa8^G470R/Wt^* mutants with their paralytic phenotype represent a useful and novel model of intermediate SMA.

In summary, this study cements the relevance of the synaptic chaperone, Hspa8, and its associated biochemical pathways to SMA biology and neuromuscular health. Modulating these pathways by either directly targeting Hspa8 or via linked factors could be broadly useful for the treatment of human neuromuscular disease.

## Acknowledgements

We thank members of the Monani lab for comments and suggestions. We are also very appreciative of D.C. De Vivo, A.H.M. Burghes and A.P. Lieberman for critically reading this manuscript and providing feedback. Finally, Manuel de Miguel and Marcos Ortega Medina are recognized for providing technical assistance and use of their histological facilities. **Funding:** This study was supported by grants from AFM-France, Cure SMA, the Muscular Dystrophy Association (MDA 1064312), the Hope for Children Research Foundation and NIH (R01 NS123292) to U.R.M., the MCIN/AEI/10.13039/501100011033 (PID2023-150602NB-100) to L.T., FPU22/02843 to A.F-M., and MDA (10.55762/pc.gr.157042) and the Canadian Institutes of Health Research (PJT-186300) to R.K. **Author contributions:** Conceptualization: Y-R.H, L.T., U.R.M.; Methodology: Y-R.H, A.F-M.; Investigation: Y-R.H, A.F-M.; Supervision: R.K., L.T., U.R.M.; Writing: Y-R.H. & U.R.M. with input from all co-authors. **Competing interests:** U.R.M. is an inventor on a patent application filed by Columbia University on the use of Hspa8 to treat neurodegenerative disease.

## Methods

### Animals

All mouse work was performed in accordance with the NIH Guidelines on the Care and Use of Lab Animals, complied with all ethical regulations, and was approved by the IACUC of Columbia University (Protocol Number AC-AABE8550). Mice were housed in a controlled environment on a 12h light/12h dark cycle with food and water, and all efforts were made to minimize suffering. *Smn^2B/2B^* and *Hspa8^G470R^* mice on the FVB/N genetic background were previously described^14,49^. *Smn^2B/2B^*; *Hspa8^G470R/G470R^* mice were crossed with *Smn^+/−^* homozygous for the G470R variant or bearing just WT Hspa8 to obtain mutants with one or two copies of the modifier. *Smn^2B/2B^* x *Smn^+/−^* crosses generated *Smn^2B/−^* mice. *Smn^2B/+^* mice served as controls in our experiments. Genotyping was performed by PCR on tail DNA using primers listed in Table S2. Mouse work in the Tabares lab was conducted in accordance with the European Council Directive for the Care and Use of Laboratory Animals and was approved by the Animal Care and Ethics Committee of the University of Seville. Righting ability of the various mouse cohorts was assessed as previously described^50^. Briefly, righting reflex was assessed daily from P5 to P10 by placing the mouse on its back and measuring the time it took to turn upright on its four paws (righting time). For each testing session, the assay was performed thrice and the mean reported. Grip strength was performed using a force transducer (Bioseb #GT3) by positioning the animals on a wire grid and gently pulling the subject horizontally along the grid. Each subject was assessed 3 – 5 times with 1min. rest intervals between tests and means reported.

### Transcript and protein assessments

Total RNA was isolated using TRIzol reagent (Invitrogen) according to the manufacturer’s instructions and subsequently reversed transcribed into cDNA with the SuperScript™ III First-Strand Synthesis System (Invitrogen). Quantitative PCR was performed in triplicate on a CFX96 Real-Time System (Bio-Rad) using the SsoAdvanced Universal SYBR Green Supermix (Bio-Rad) and raw values normalized to endogenous Gapdh or β-Actin mRNA levels; for primer sequences, see Table S2. For immunoblot analysis, tissue or cell lysates from Mouse Embryonic Fibroblasts (MEFs) were used. Briefly, lysates were made with modified RIPA buffer (50 mM Tris-HCl pH 8.0, 150 mM NaCl, 1% NP-40, 0.5% sodium deoxycholate, 0.1% SDS and 1X protease/phosphatase inhibitors cocktail (Roche) and boiled at 95°C for 5 minutes in 1X SDS sample buffer. For tissue-derived protein, tissue was homogenized and incubated on ice for 20 min. Lysates were then centrifuged for 30 min at 14,000g to remove the insoluble debris. Protein amounts were assessed using the BCA protein assay kit (Thermofisher) and concentrations estimated on an Infinite F50 Plus plate reader supplied with Magellan data analysis software (Tecan). Western blotting was conducted by separating proteins (10-50μg) on 10% or 12% SDS-PAGE gels and then transferring the proteins to a PVDF membrane (Millipore Inc.) as previously described^51^. Blots were blocked in 1X TBST (Tris-buffered saline with 0.1% Tween 20) containing 5% nonfat milk, and primary antibody used to probe blots for 1hr at room temperature or for 18hrs at 4°C according to the supplier’s recommendations. Following incubation with primary antibody membranes were washed (1X TBST) 3 times with agitation for 5 minutes at room temperature and then incubated with appropriate secondary antibodies for 1hr at room temperature. Antibodies used for the western blotting are as follows: SMN (1:1000, BD Biosciences, Cat#610647), p62 (1:1000, Cell Signaling, Cat#5114), β-tubulin (1:1000, Santa Cruz, Cat#SC53140), α-tubulin (1:1000, Santa Cruz, Cat#SC-398103), β-actin (1:1000, Santa Cruz, Cat#SC47778), Goat anti-Rabbit IgG-HRP (1:10,000, Jackson ImmunoResearch, Cat#111-035-003) and Goat anti-Mouse IgG-HRP (1:10,000, Jackson ImmunoResearch, Cat# 115-035-003). Protein bands were visualized using the ECL kit (Bio-Rad Inc.) and images captured on a ChemiDoc Imaging System (Bio-Rad Inc.). Band intensities were assessed using the ImageJ software (NIH, Bethesda, MD, USA) or Image Lab (Bio-Rad Inc.).

### Plasmids and constructs

The SMN-FLAG and Hspa8-Myc expression vectors were generated by inserting the full-length murine *Smn* and murine *Hspa8* cDNAs into the Ecor1-Xho1 and Kpn1-Xho1 sites of pcDNA3.1-FLAG and pcDNA3.1-Myc respectively. The full-length constructs were used as templates to generate all deletion, truncation and substitution mutants (Gene Synthesis system, GeneScript). The various constructs used are as follows: SMN(FL)-FLAG, SMN^ΔG^-FLAG, SMN^ΔT^-FLAG, SMN^ΔP^-FLAG, SMN^ΔY^-FLAG, SMN_N-FLAG, Hspa8^WT^-Myc, Hspa8^G470R^-Myc, Hspa8_NBD-Myc, Hspa8_SBD-Myc and Hspa8^ΔLid^-Myc. The amino acid boundaries corresponding to each deletion or truncation mutant are detailed in Fig. S4. All constructs were verified by DNA sequencing.

### Cell cultures, transfections and co-immunoprecipitation studies

Mouse embryonic fibroblasts were obtained from either WT or Hspa8^G470R/G470R^ E12.5-day-old embryos. Cells were cultured in DMEM (5% CO_2_ at 37°C) supplemented with 10% fetal bovine serum, 100 units/ml penicillin and 100μg/ml streptomycin. To measure the half-life of SMN protein and mRNA, cells were treated with 80μg/ml of cycloheximide and 7ug/ml actinomycin D (Sigma-Aldrich), respectively for the indicated periods of time. To inhibit lysosomal activity, cells were treated with 100μM leupeptin (Fisher BioReagents, Thermo Fisher Scientific) and 20mM ammonium chloride (Sigma-Aldrich) for 6 hours with, or without 80ug/ml of cycloheximide.

To identify domains in Hspa8 and SMN that mediate their interactions with each other, Myc and FLAG-tagged versions of these cDNAs were co-transfected [Lipofectamine 3000 (Life Technologies, Thermo Fisher Scientific)] into and over-expressed in HEK293 cells. Twenty-four hours later, cells were washed once with ice-cold PBS then harvested in NP-40 lysis buffer containing 50mM Tris-HCl pH 7.5, 150mM NaCl, 1% NP-40, 10% glycerol and 1X protease/phosphatase inhibitors cocktail (Roche). The resulting lysates were clarified by centrifugation at 16,000g for 20 min at 4°C, then pre-cleared by incubating with 20μl of Protein A/G Plus Agarose (Thermofisher Inc.) for 1hr. Next, SMN-FLAG was immunoprecipitated using 20μl of anti-DYKDDDDK IP Resin (GenScript, Cat#L00425) by end-over-end rotation (8hrs at 4°C). The immunoprecipitated proteins were washed (3X) with ice-cold lysis buffer and following the final wash, the supernatant was removed and the beads resuspended in 1X SDS sample, boiled for 5mins., resolved by SDS-PAGE and then subjected to immunoblot analysis as described above. Blots were probed using anti-DYKDDDDK (1:1000, GenScript, Cat#A00187), anti-MYC (1:1000, Cell Signaling, Cat#2276), anti-Bag3 (1:1000, Proteintech, Cat#10599-1-AP) and/or anti-Hspb8 (1:1000, Proteintech, Cat# 15287-1-AP) antibodies as indicated in Fig. 4. All immunoblots depicted are representatives of at least three experiments that demonstrated similar results. Note that detection of SMN-p62 and SMN-Hspb8 complexes was enhanced by incubating SMN-FLAG immunoprecipitate with ATPγS, a modified form of ATP relatively slow to undergo hydrolysis and therefore suitable for identifying ATP-bound intermediate CASA complexes. For this, following the final wash of immunoprecipitated SMN-FLAG complexes, proteins were resuspended in nucleotide buffer (25mM HEPES pH 7.4, 100mM KCl, 5mM MgCl₂, 0.1% NP-40) containing 1mM ATPγS and 5mM MgCl_2_. Bead suspensions were incubated for 5min on ice with occasional gentle mixing. Following incubation, beads were briefly centrifuged, washed once with NP-40 wash buffer to remove excess nucleotide, supernatant removed and beads boiled in 2X SDS sample buffer for 5 min. Eluates were analyzed by SDS–PAGE and immunoblotting as described above.

### Anatomical studies

Motor neuron, NMJ and muscle histology were essentially carried out as previously described^14,50^ and detailed below.

#### Motor neurons

Mice were euthanized, subjected to transcardial perfusion [4% PFA in PBS (pH 7.4)] and lumbar spinal cord (L1-L5) extracted. The tissue was then post-fixed in the same fixative, cryo-protected initially in 20% and then 30% sucrose before embedding it in Tissue-Tek Optimal Cutting Temperature (OCT) compound (Fisher) for cryostat sections. All embedded tissue was frozen on dry ice and stored at -80°C until processing. Cryosections (20μm) were collected onto Superfrost Plus glass slides (Fisher) using a CM3050S cryostat (Leica). The sections were washed once with 1X PBS for 5 minutes to remove OCT, overlaid with 4% PFA for 10 min, washed again (1X PBS) and then permeabilized with 0.5% Triton X-100 (5 min.). To stain the motor neurons, sections were placed in blocking buffer (5% normal donkey serum, 3% BSA, 0.1% Triton X-100 in PBS) for 1h, then incubated (4°C, overnight) with anti-ChAT primary antibody (1:100, Millipore, Cat#AB144P) diluted in blocking buffer, following which they were washed (4 X 15 min) in 1X PBST. The following day, the sections were incubated with secondary antibody, Alexa Fluor-594 conjugated donkey anti-goat IgG (1:1000, Invitrogen), for 2h. After a second round of washing (4 X15 min.) in 1X PBST, the sections were mounted in Vectashield (Vector Labs, Burlington, VT, USA) containing DAPI. Motor neurons were visualized either or a Nikon 80i fluorescent microscope (Nikon) or a Leica TCS SP8 laser scanning confocal microscope (Leica). ChAT-positive motor neurons soma in the ventral horns were enumerated between L1 and L5 regions of the spinal cord; sections were analyzed at an effective spacing of ∼50μm to avoid counting soma twice.

#### NMJs

NMJs of triceps brachii or gastrocnemius muscles were stained as previously described^49^. Briefly, muscle tissue was fixed and permeabilized with 100% methanol (10 min, −20°C), then teased longitudinally into small bundles under a dissecting microscope before incubating (1h) with blocking buffer (5% normal serum, 0.1% Triton X-100 in PBS). It was then sequentially incubated for overnight periods at 4°C with an anti-neurofilament antibody (1:1000, Millipore), followed by Alexa Fluor-488 conjugated donkey anti-rabbit IgG secondary antibody (1:1000, Invitrogen) and rhodamine-α-bungarotoxin (1:1000, Invitrogen). Each incubation was followed by a washing step (4 X in 1X PBST for 15 min). Tissues were then mounted in anti-fade medium (Vector Labs), and NMJs imaged using a Leica SP8 confocal microscope equipped with LAS X software (v1.9.0.13747). NMJs exhibiting no innervation or, <50% overlap between NF and α-bungarotoxin–label, were classified as denervated. Characterization and quantification of innervated NMJs, defective terminals, and motor endplates was conducted as previously described^14,50^. All images were analyzed off-line using the Leica LAS X software (v1.9.0.13747).

#### Muscles

Muscle was flash-frozen in isopentane chilled with liquid nitrogen, then embedded in Tissue-Tek OCT medium, and 12-µm transverse cryosections cut on a Leica cryostat (Leica, Deerfield). Sections were stained with hematoxylin and eosin (Sigma-Aldrich) and mounted using Cytoseal XYL (Fisher Scientific). Muscle morphology, fiber size, and fiber number were evaluated using a Nikon Eclipse 80i fluorescence microscope equipped with a SPOT 4.5 camera (Diagnostic Instruments, Sterling Heights, MI, USA) and analyzed with ImageJ software (NIH, Bethesda, MD, USA). Fiber size measurements were obtained from ≥100 fibers per sample.

#### Liver

Livers were fixed in 4% paraformaldehyde for 48-72 h at 4 °C and then processed at the Department of Normal and Pathological Cytology and Histology at the University of Seville. All the samples were dehydrated, cleared with xylol, and embedded in paraffin wax using a Spin Tissue Processor STP 120 (Myr). Paraffin block tissues were cut with a microtome (HM 310, Thermo Scientific) at 4 µm thickness. Sections were stained with Haematoxylin and Eosin (H&E) using standard methods and assessed by light microscopy (Zeiss Axiovert 40C, x10, x40 objectives).

### AAV-mediated Hspa8^G470R^ treatment

Mice were administered AAV-PHP.eB-Hspa8^G470R^ (6X10^10^ GC per pup in a volume of 2μl) at PND0 via the cerebral ventricles. Briefly, pups were cryo-anesthetized on a paper towel on wet ice for 3 mins until no pedal withdrawal reflex was observed. ICV injections were conducted using a 1-mL insulin syringe with a pulled glass capillary needle tip (1μm diameter). Vectors were injected into the left and right hemisphere 2mm lateral to the midline, midway between bregma and lambda, and to a depth of 1mm below the surface of the skull. Following injection, pups were placed on a warming blanket and returned to the dam in their own cage once they were visibly active. Pups were monitored every day following virus delivery and were weaned at 21days.

### Acute neuromuscular preparations

Mice were euthanized with 100% CO₂ and the TVA muscle dissected as previously described^10^. Preparations were continuously perfused with a solution containing (in mM): 135 NaCl, 4 KCl, 2 CaCl₂, 1 MgCl₂, 15 NaHCO₃, 0.33 NaH₂PO₄, and 10 glucose. The solution was equilibrated with a gas mixture of 95% O₂ and 5% CO₂.

### Electrophysiology

The motor nerve was stimulated using a suction electrode with square wave pulses of 0.15 ms duration and 2–10 mV amplitude. Intracellular recordings from single muscle fibers near motor endplates were obtained using glass microelectrodes (10–20 MΩ) filled with 3 M KCl and connected to an intracellular amplifier (TEC-05X; NPI Electronic, Germany). Evoked endplate potentials (EPPs) and miniature EPPs (mEPPs) were recorded at room temperature (22–23°C), as previously described^38^. Muscle contractions were blocked by adding 2 μM μ-conotoxin GIIIB (Alomone Labs, C-270, Israel) a selective inhibitor of voltage-gated sodium channels in skeletal muscle, to the bath. Recordings were sampled at 20 kHz. EPP and mEPP amplitudes were normalized to a resting membrane potential of −70 mV, and EPPs were corrected for nonlinear summation^52^. The quantal content *(m)* was calculated using the direct method:

*m*=Average mEPP amplitude/Average EPP amplitude. During high-frequency stimulation trains, *m* was calculated as the ratio between each EPP and the mean mEPP amplitude under each experimental condition. To estimate the readily releasable pool (RRP) size, *m* values during the train were plotted over time and fitted to a sequential model as described^37^. The model assumes that quanta released upon stimulation originate from the RRP, which is depleted exponentially. Recruitment begins after the first stimulus and rises sigmoidally to a plateau as the RRP depletes. The curve of quantal content over time, *m(t)*, was fitted to the following function:

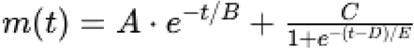

where, A = Initial quantal content, B = Time constant of RRP depletion, C = Plateau amplitude, D = Vesicle recruitment half-time, and E = Steepness of the recruitment curve. Note: The sigmoid component was constrained to begin at zero. Integration of the first exponential component provided an estimate of the RRP size (also see Fig. S7).

### Statistical analyses

Sample sizes for the various experiment are detailed in the figure legends. Kaplan-Meier survival curves were assessed for differences using the log-rank test equivalent to the Mantel-Haenszel test. Two-tailed unpaired Student’s *t*-test or Mann-Whitney tests – when data was not normally distributed – were employed to compare pairs of cohorts with one another. One-way ANOVA followed by Tukey’s post-hoc comparison, or the Kruskal-Wallis test followed by Dunn’s post-hoc comparison – when data was not normally distributed – were used to compare three or more cohorts for statistical differences. For protein and RNA stability comparisons, AUC (Area Under Curves) were calculated and an overall *P* value assigned based on *t*-tests of the AUCs. Data are represented as mean ± SEM unless otherwise indicated. *P* < 0.05 was considered significant. Statistical analyses were performed with GraphPad Prism v9.5.1 (GraphPad Software).

## Data availability

Underlying values associated with data presented in the study may be requested and will be made available by the lead contact, Umrao R. Monani.

## Supplemental Information

**Table S1 –** Single nucleotide polymorphism (SNP) analysis of *Smn^2B/-^* breeders used to generate mutants for the study.

**Table S2 –** Key resources used for the study.

**Video S1 –** Video depicting hindlimb paralysis in *Smn^2B/-^;Hspa8^G470R/Wt^* mutants but not a littermate homozygous for the modifier *(Smn^2B/-^;Hspa8^G470R/G470R^)*.

