## Supplementary material for "Protein-stabilizing and neurotransmission-potentiating activities of a synaptic chaperone modify spinal muscular atrophy in model mice": Table S2

**Supplemental Table 2 (Key Resources)**

| REAGENT or RESOURCE | SOURCE | IDENTIFIER |
|---|---|---|
| *Antibodies* | | |
| SMN | BD biosciences | Cat#610647; RRID: AB_397973 |
| α-tubulin | Santa Cruz | Cat#SC398103; RRID: AB_2832217 |
| β-tubulin | Santa Cruz | Cat#SC53140; RRID: AB_793543 |
| β-actin | Santa Cruz | Cat#SC47778; RRID: AB_626632 |
| Hspa8 | Invitrogen | Cat#MA1-26078; RRID: AB_794523 |
| SQSTM1/p62 | Cell Signaling | Cat#5114; RRID: AB_10624872 |
| LC3B | Cell Signaling | Cat#2775; RRID: AB_915950 |
| BAG3 | Proteintech | Cat#10599-1-AP; RRID: AB_2062602 |
| Hspb8 | Proteintech | Cat#15287-1-AP; RRID:AB_2248640 |
| DYKDDDDK-Tag | GenScript | Cat#A00187; RRID:AB_1720813 |
| Myc-Tag | Cell Signaling | Cat#2276; RRID:AB_331783 |
| Goat anti-Rabbit IgG-HRP | Jackson ImmunoResearch | Cat#111-035-003; RRID: AB_2313567 |
| Goat anti-Mouse IgG-HRP | Jackson ImmunoResearch | Cat#115-035-003; RRID: AB_10015289 |
| Choline acetyltransferase (ChAT) | Millipore | Cat#AB144P; RRID: AB_2079751 |
| Neurofilament | Millipore | Cat#AB1987; RRID: AB_91201 |
| Rhodamine-α-bungarotoxin | Invitrogen | Cat#B13424; RRID: AB_2313931 |
| Alexa Fluor-488 conjugated goat anti-rabbit | Invitrogen | Cat#A-11034; RRID: AB_2576217 |

| | | |
|---|---|---|
| Alexa Fluor-594 conjugated donkey anti-goat | Invitrogen | Cat#A-11058; RRID: AB_2534105 |
| *Chemicals and reagents* | | |
| cycloheximide | Sigma-Aldrich | Cat#C7698 |
| actinomycin D | Sigma-Aldrich | Cat#A9415 |
| ammonium chloride | Sigma-Aldrich | Cat#A9434 |
| leupeptin | Sigma-Aldrich | Cat#L5793 |
| ATPγS | Sigma-Aldrich | Cat#A1388 |
| Lipofectamine™ 3000 Transfection Reagent | Invitrogen | Cat#L3000015 |
| Pierce™ Protein A/G Plus Agarose | Thermo Scientific | Cat#20423 |
| anti-DYKDDDDK IP Resin | GenScript | Cat#L00425 |
| *Software and Algorithms* | | |
| GraphPad Prism | Graph Pad Software | RRID: SCR_002798 |
| ImageJ | NIH | RRID: SCR_003070 |
| Image Lab | Bio-Rad | RRID:SCR_014210 |
| Leica LAS X | Leica | RRID: SCR_013673 |
| *PCR primers for genotyping* | | |
| $Smn^{WT}$ | This paper | F 5' GGTGCTAGGATCTCTGTGTTCGTGC 3'<br>R 5' GGTCCCACCACCTAAGAAAGC 3' |
| $Smn^{KO}$ | This paper | F 5' GCTGCCCATCCACCCTCTG 3'<br>R 5' GGTCCCACCACCTAAGAAAGC 3' |
| $Smn^{2B}$ | This paper | F 5' AACTCCGGGTCCTCCTTCCT 3'<br>R 5' TTTGGCAGACTTTAGCAGGGC 3' |
| $Hspa8^{WT}$ | This paper | F 5' ATCCCTCCAGCACCACTTG 3'<br>R 5' GCTTCTCATCCTCAGCCTTG 3' |
| $Hspa8^{G470R}$ | This paper | F 5' CACCAGAAATGCTGTGTTGG 3'<br>R 5' AGTAACCTCAATCTGAGGGACTCG 3' |

| PCR primers for RT-qPCR | | |
|---|---|---|
| β-actin | This paper | F 5’ TGTTACCAACTGGGACGACA 3’<br>R 5’ GGGGTGTTGAAGGTCTCAAA 3’ |
| Gapdh | This paper | F 5’ TGGAGAAACCTGCCAAGTATGA 3’<br>R 5’ GCCGTATTCATTGTCATACCAGG 3’ |
| Smn-FL | This paper | F 5’ CACCACCTCCCATATGTCCAGATT 3’<br>R 5’ GAATGTGAGCACCTTCCTTCTTT 3’ |
| Smn-Δ7 | This paper | F 5’ GGACCACCAATAATCCCGCC 3’<br>R 5’CATATAGAAGATAGAAAAAAACAGTAC AATGAAC 3’ |
| Smn-Total | This paper | F 5’ GAAGATACGGTGCTGTTCCG 3’<br>R 5’ AGCATGCTTAAAGGAAGCCA 3’ |
