## Supplemental Figures for "Protein-stabilizing and neurotransmission-potentiating activities of a synaptic chaperone modify spinal muscular atrophy in model mice"

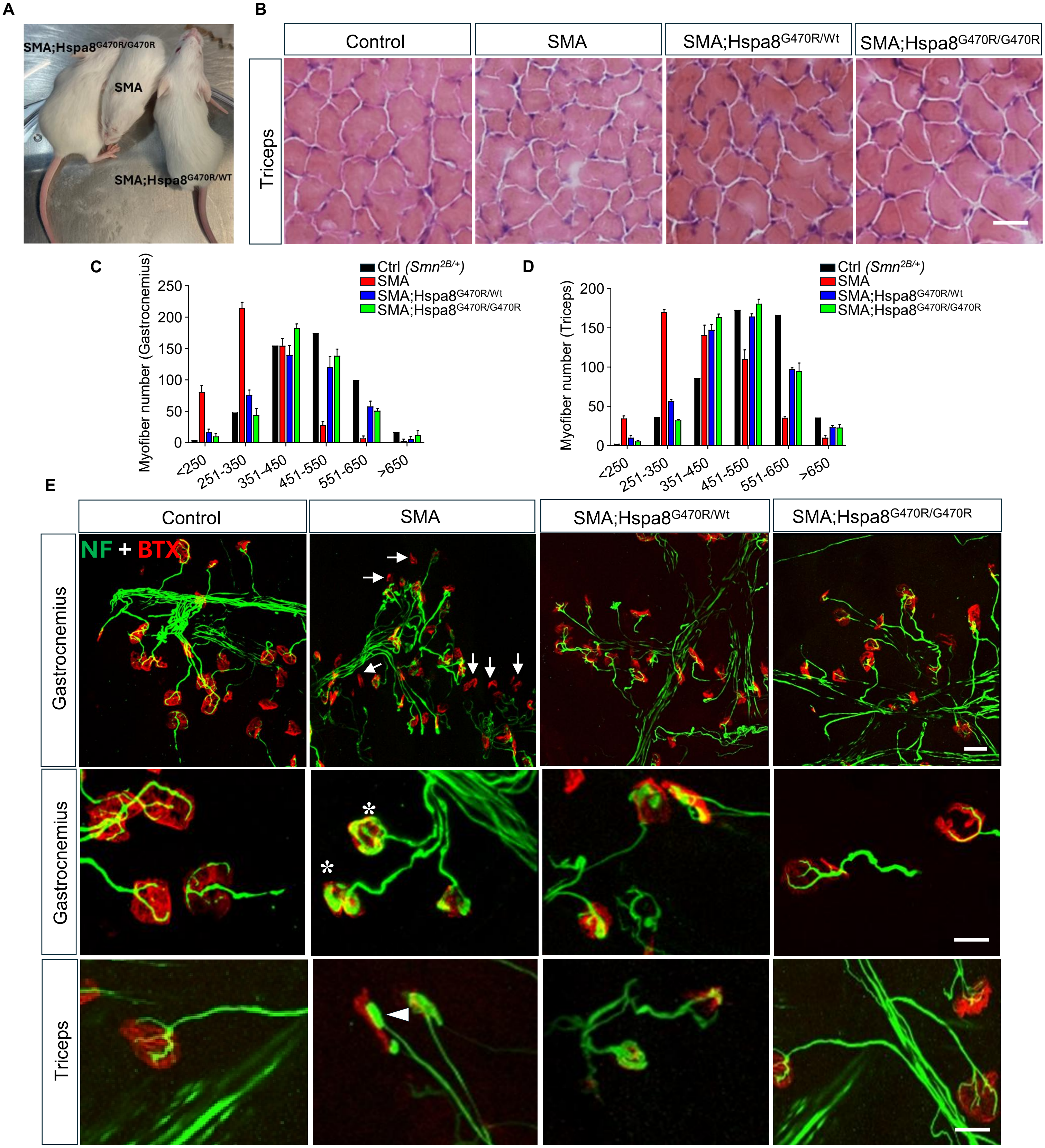


**Supplemental Figure 1 – The Hspa8$^{G470R}$ variant improves health and ameliorates neuromuscular pathology in SMA model mice. (A)** Representative PND15 SMA mutants with or without the disease modifier. Note smaller size of mutant lacking the modifier. **(B)** H&E-stained sections from the triceps muscles of controls and mutants with or without the SMA modifier. Scale bar – 25μm. **(C)** Histograms showing distribution of myofiber areas from **(C)** the gastrocnemius and **(D)** the triceps of the various cohorts of mice. Note rightward shift and normalization of fiber size in mutants with the modifier. **(E)** Immunostains of NMJs from each of the four groups of mice. Highlighted are denervated NMJs (arrows), endplates exhibiting nerve terminals engorged with NF protein (asterisks) and axons terminating as end-bulbs in *Smn$^{2B/-}$* mice. These pathologies are lessened or abolished in the presence of the modifier. Scale bars – 20μm. Data: mean ± SEM

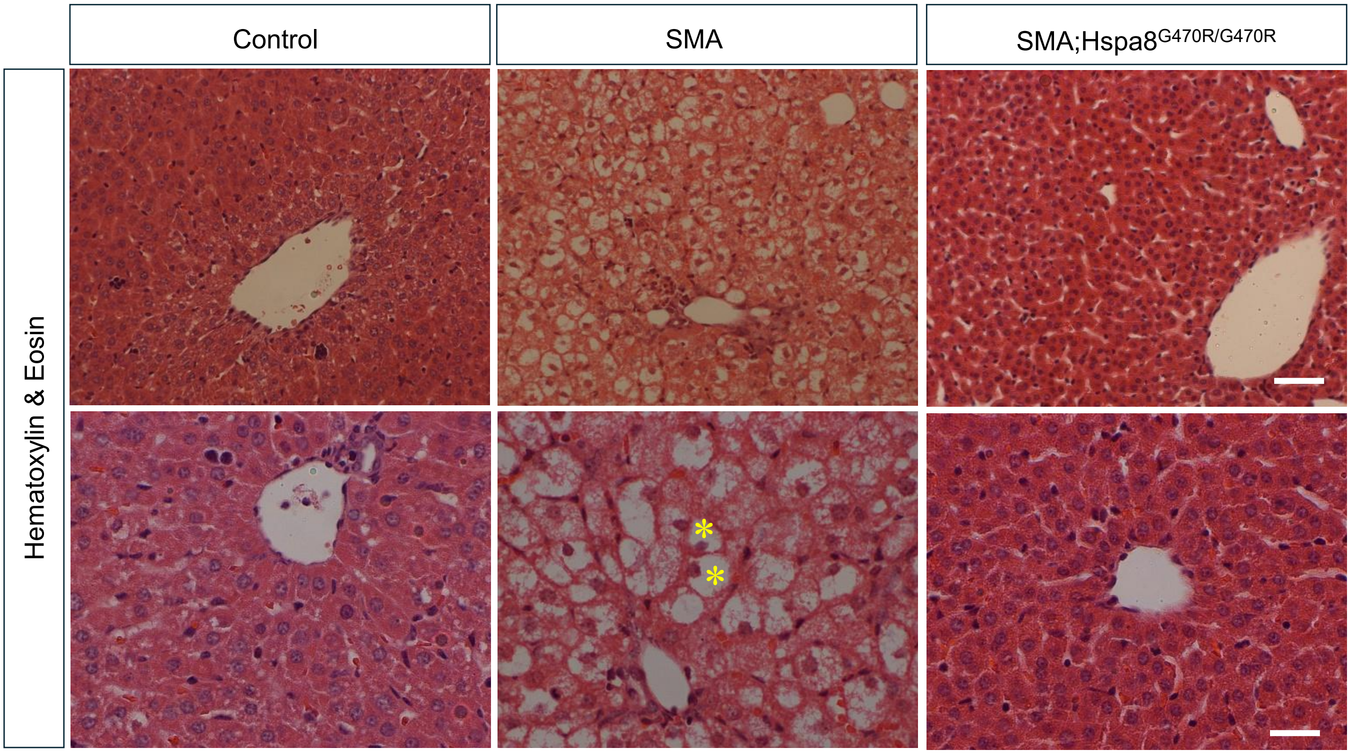


**Supplemental Figure 2 – The Hspa8$^{G470R}$ variant abolishes liver pathology in SMA model mice.** Representative histological sections of liver tissue stained with hematoxylin-eosin of a PND15 healthy control and age-matched SMA mutants devoid of the Hspa8$^{G470R}$ variant or homozygous for it. Note the microvesicular steatosis and foamy hepatocytes (asterisks) in the *Smn*$^{2B/-}$ mouse liver, which is effectively corrected by the modifier, restoring normal hepatocyte architecture. Scale bars – 30μm and 15μm, upper and lower panels respectively.

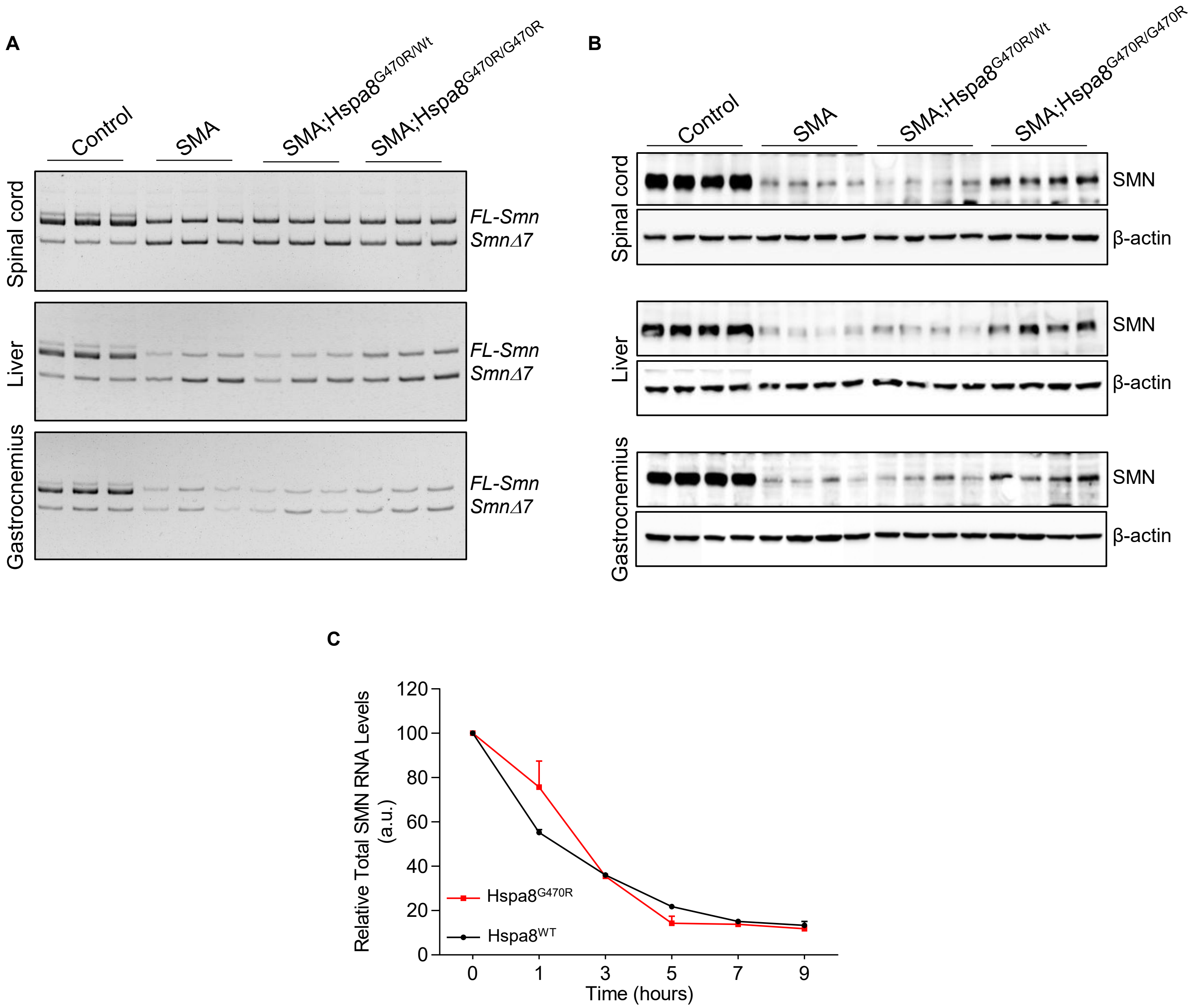


**Supplemental Figure 3 – The Hspa8$^{G470R}$ variant modulates the SMN protein but not transcript levels. (A)** Representative polyacrylamide gel electrophoresis of the FL and Δ7 RNA isoforms of the $Smn^{2B}$ allele in diverse tissues of controls and SMA mutants with or without the Hspa8$^{G470R}$ disease modifier. Note equivalent levels of the FL transcript in all SMA mutants. **(B)** Representative western blots of SMN protein in different tissues of controls and mutants expressing either the WT or variant form of Hspa8. SMN increase is not noticeable in mutants with one variant allele but is observed in mutants with two variant alleles relative to SMA mice expressing WT Hspa8. **(C)** Graphical representation of murine *Smn* RNA decay in MEFs either WT for Hspa8 or expressing the variant form of it, following treatment with Actinomycin D to halt transcription. Note: Curve comparison by quantifying areas under curves followed by a *t* test failed to reveal a difference between the two genotypes; results compiled from 3 independent experiments. Data: mean ± SEM

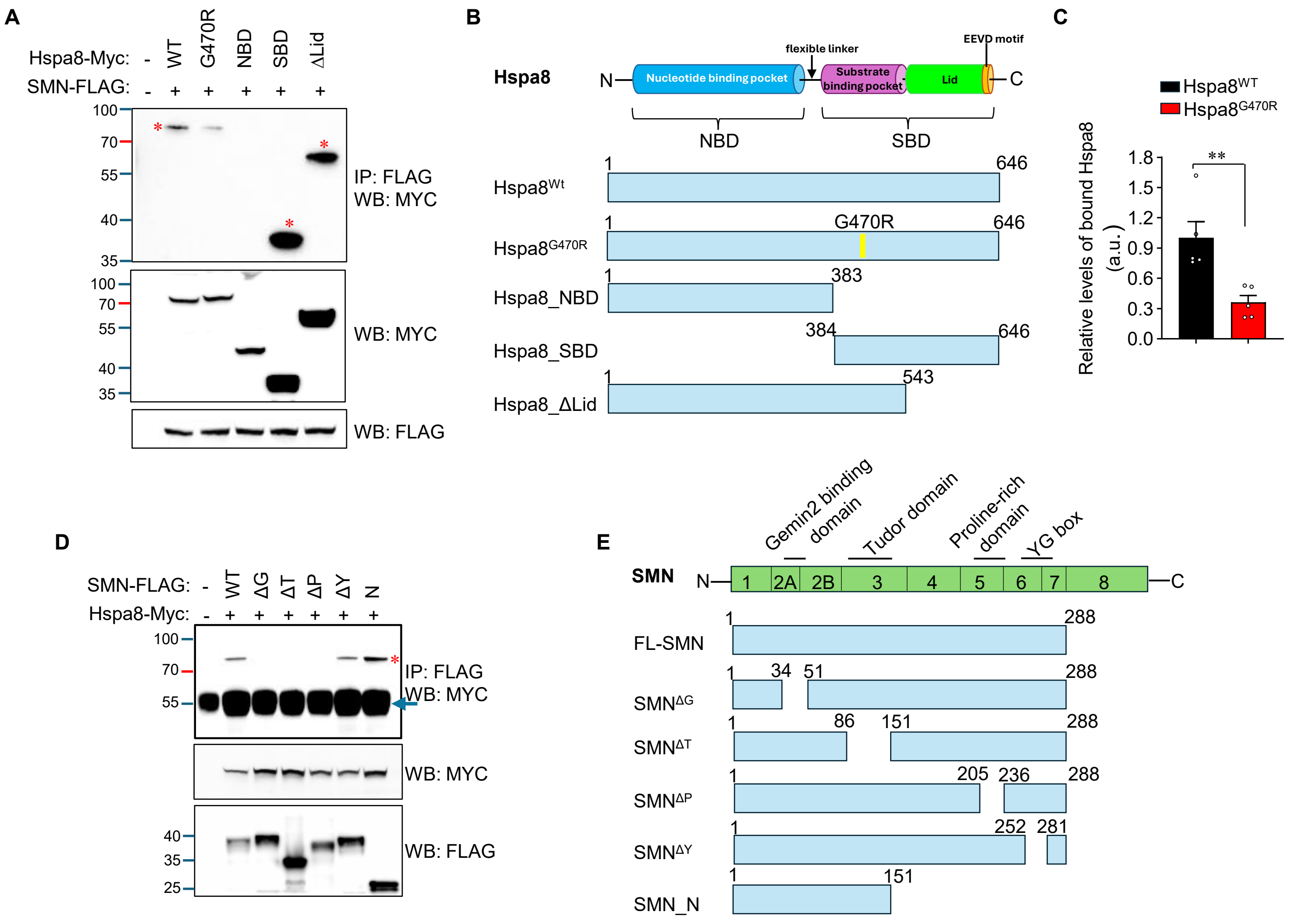


**Supplemental Figure 4 – The substrate-binding and Tudor domains of Hspa8 and SMN respectively mediate their interaction. (A)** Representative blot following co-IP of various Hspa8-Myc deletion constructs with full-length SMA-FLAG. Note reduced binding of the G470R variant versus Hspa8$^{WT}$ protein to SMN; asterisks denote bands of interest **(B)** Cartoon of the various domains of Hspa8 and the respective chaperone constructs employed for the co-IP experiment in panel A. **(C)** Quantified relative amounts of the WT or the G470R variant of Hspa8 bound to SMN in co-IP experiments represented by panel A. **(D)** Representative blot following co-IP of full-length Hspa8-Myc protein with protein derived from various SMN-FLAG deletion constructs; asterisk denotes band of interest. **(E)** Cartoon of the various domains of SMN and the respective SMN constructs employed for the co-IP experiment in panel D.

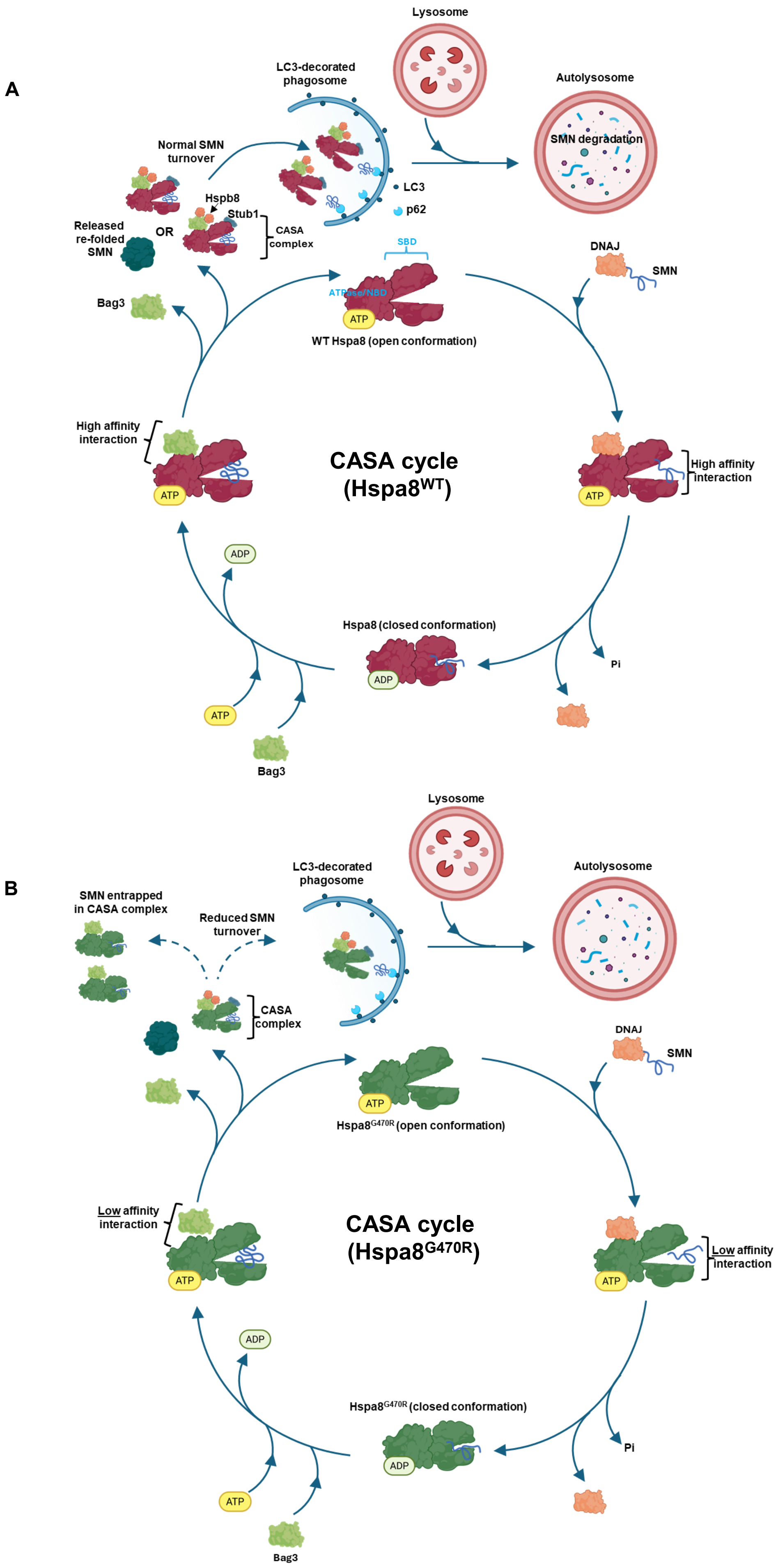
A
Lysosome
LC3-decorated phagosome
Autolysosome
SMN degradation
Normal SMN turnover
Hspb8
Stub1
CASA complex
LC3
p62
Released re-folded SMN
OR
SBD
ATPase/NBD
ATP
WT Hspa8 (open conformation)
DNAJ
SMN
Bag3
High affinity interaction
CASA cycle (Hspa8$^{WT}$)
High affinity interaction
ADP
Hspa8 (closed conformation)
Pi
B
Lysosome
LC3-decorated phagosome
Autolysosome
SMN entrapped in CASA complex
Reduced SMN turnover
CASA complex
DNAJ
SMN
Hspa8$^{G470R}$ (open conformation)
Low affinity interaction
CASA cycle (Hspa8$^{G470R}$)
Low affinity interaction
ADP
Hspa8$^{G470R}$ (closed conformation)
Pi
ATP
Bag3

**Supplemental Figure 5 – Key steps in chaperone-assisted selective autophagy.** Schematic of Hspa8-mediated chaperone-assisted selective autophagy of SMN in the presence of **(A)** WT Hspa8 (upper panel) and **(B)** Hspa8$^{G470R}$ (lower panel). Highlighted in the lower panel is the weakened interaction identified in this study (also see Fig. 4) between SMN and the G470R variant on the one hand and between Bag3 and the G470R variant on the other. Also depicted in the lower panel is a representation of the entrapment of SMN in the multi-protein CASA complex and its reduced delivery to the p62 adaptor for degradation in lysosomes. Note: Some aspects of the cycle are generic, and it is unclear if there is a specific DNAJ protein responsible for delivery of SMN to Hspa8. Also note: NBD – nucleotide binding domain; SBD – substrate binding domain.

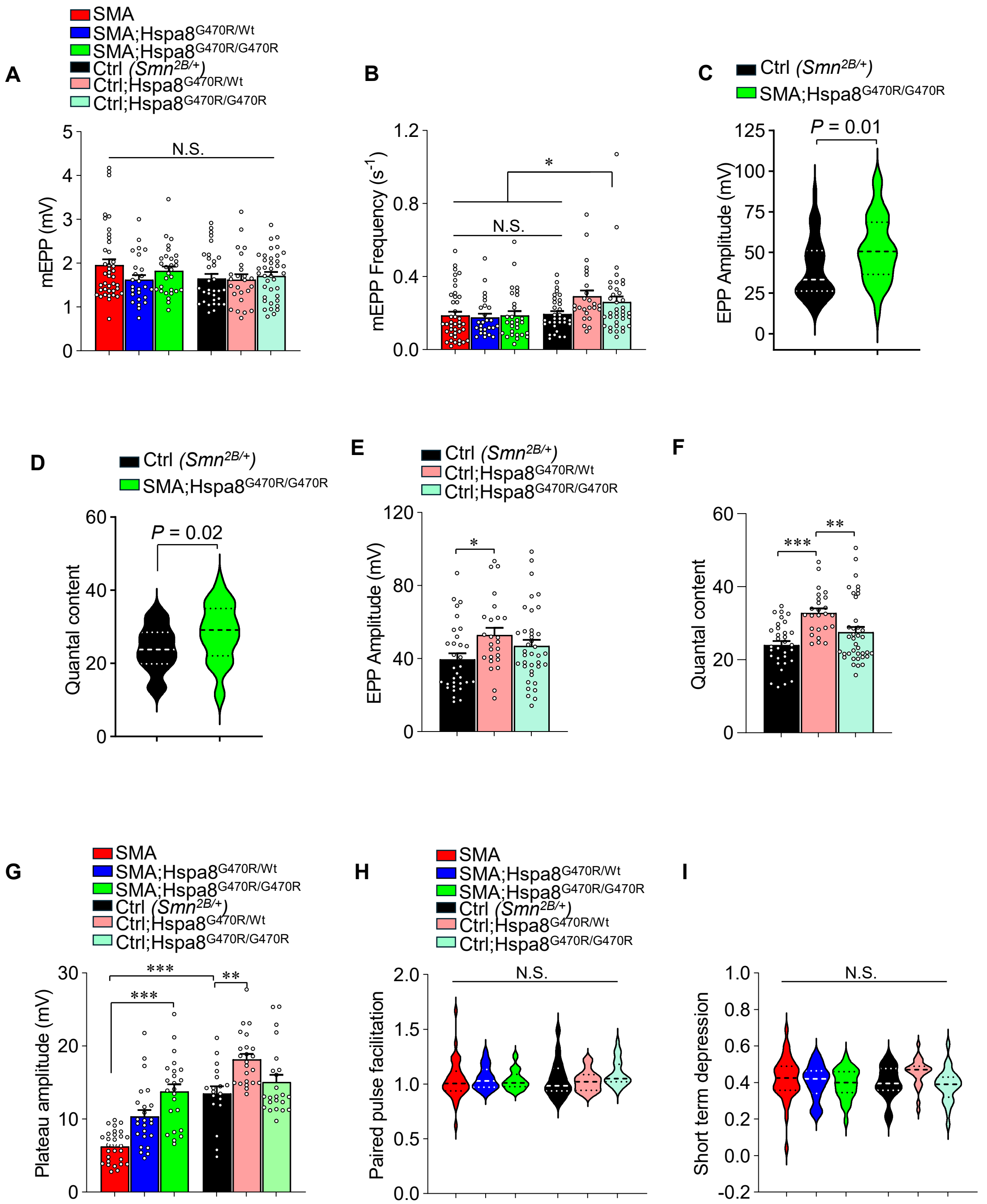


**Supplemental Figure 6 – $Hspa8^{G470R}$ enhances synaptic efficacy without raising SMN correspondingly.** The G470R chaperone variant alters neither **(A)** mEPP amplitude nor **(B)** mEPP frequency in PND15 SMA mutants. Graphical representations of head-to-head comparisons of **(C)** evoked potentials and **(D)** quantal content between healthy controls and SMA mutants homozygous for $Hspa8^{G470R}$. Each measure was significantly higher (*, $P < 0.05$, *t* test) in the mutants despite expressing lower SMN. $Hspa8^{G470R}$ raised **(E)** EPPs and **(F)** quantal content significantly in healthy controls. Note: *, **, ***, $P < 0.05$, $P < 0.01$ and $P < 0.001$ respectively, one-way ANOVA (panel E) and Kruskal-Wallis test (panel F). **(G)** Quantification of plateau amplitudes at NMJs of the various cohorts of mice subjected to high-frequency stimulation trains illustrate the propensity of $Hspa8^{G470R}$ to enhance neurotransmission. Note: **, ***, $P < 0.01$ and $P < 0.001$ respectively, Kruskal-Wallis test. $Hspa8^{G470R}$ altered neither **(H)** paired-pulse facilitation (EPP2/EPP1) nor **(I)** short-term depression [1 – (mean EPP30 – 40)/EPP1] significantly at NMJs of mutants and controls; Kruskal-Wallis test (panel H) and one-way ANOVA (panel I). Note: All analyses depicted here are based on means calculated from n ≥ 25 NMJs from N = 3 – 4 mice of each genotype. Data: mean ± SEM

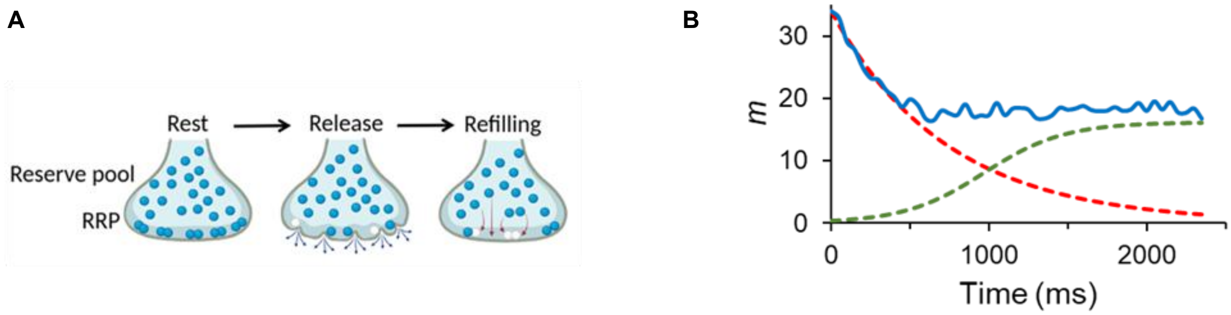


**Supplemental Figure 7 – Neurotransmitter release at the mammalian NMJ. (A)** Cartoon depicting the localization, mobilization and refilling of the various pools of synaptic vesicles at the mouse NMJ prior to, during and following neurotransmitter release. At rest, synaptic vesicles are docked at active zones forming the RRP. Upon action potential arrival, vesicles are released, recycled via endocytosis, and replaced by reserve pool vesicles to replenish the RRP. **(B)** The kinetic model of vesicle mobilization, depletion and replenishment adapted from Ruiz et al, 2011 and used to estimate RRP in the study. Calculated quantal content (*m*, blue line) is plotted over time, alongside the predicted depletion of the RRP (red dashed line) and its replenishment (green dashed line).

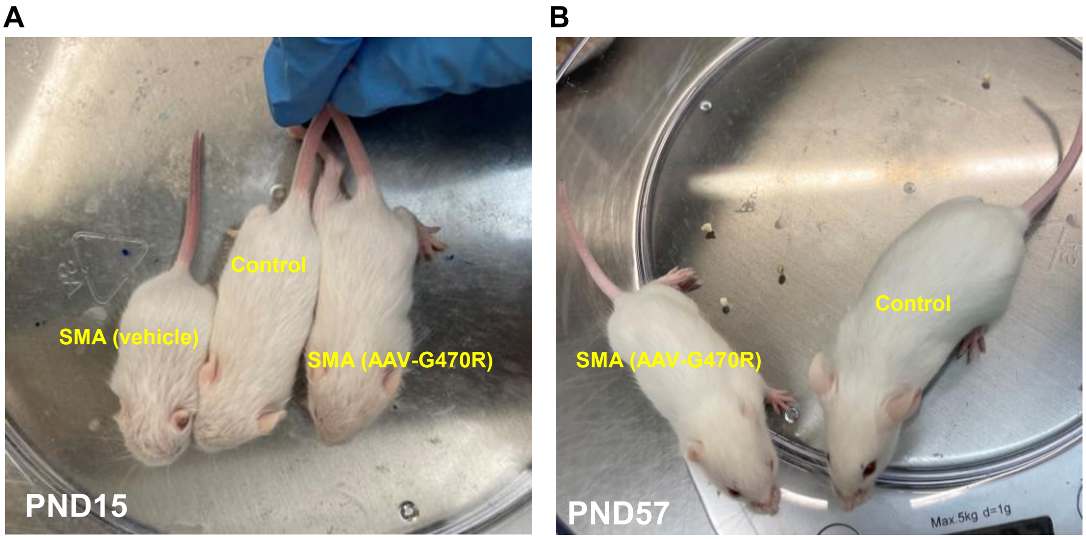


**Supplemental Figure 8 – AAV-PHP.eB-Hspa8$^{G470R}$ delivered postnatally improves the health of SMA model mice. (A)** Panel depicts a representative PND15 healthy control *(Smn$^{2B/+}$)* animal and age-matched SMA mutants *(Smn$^{2B/-}$)* that were administered either AAV-PHP.eB-Hspa8$^{G470R}$ or vehicle at birth. Note larger size and less disheveled appearance of the mutant that received a dose of the Hspa8 modifier. **(B)** Depiction of the virus-treated mutant in panel A and its control littermate at PND57.

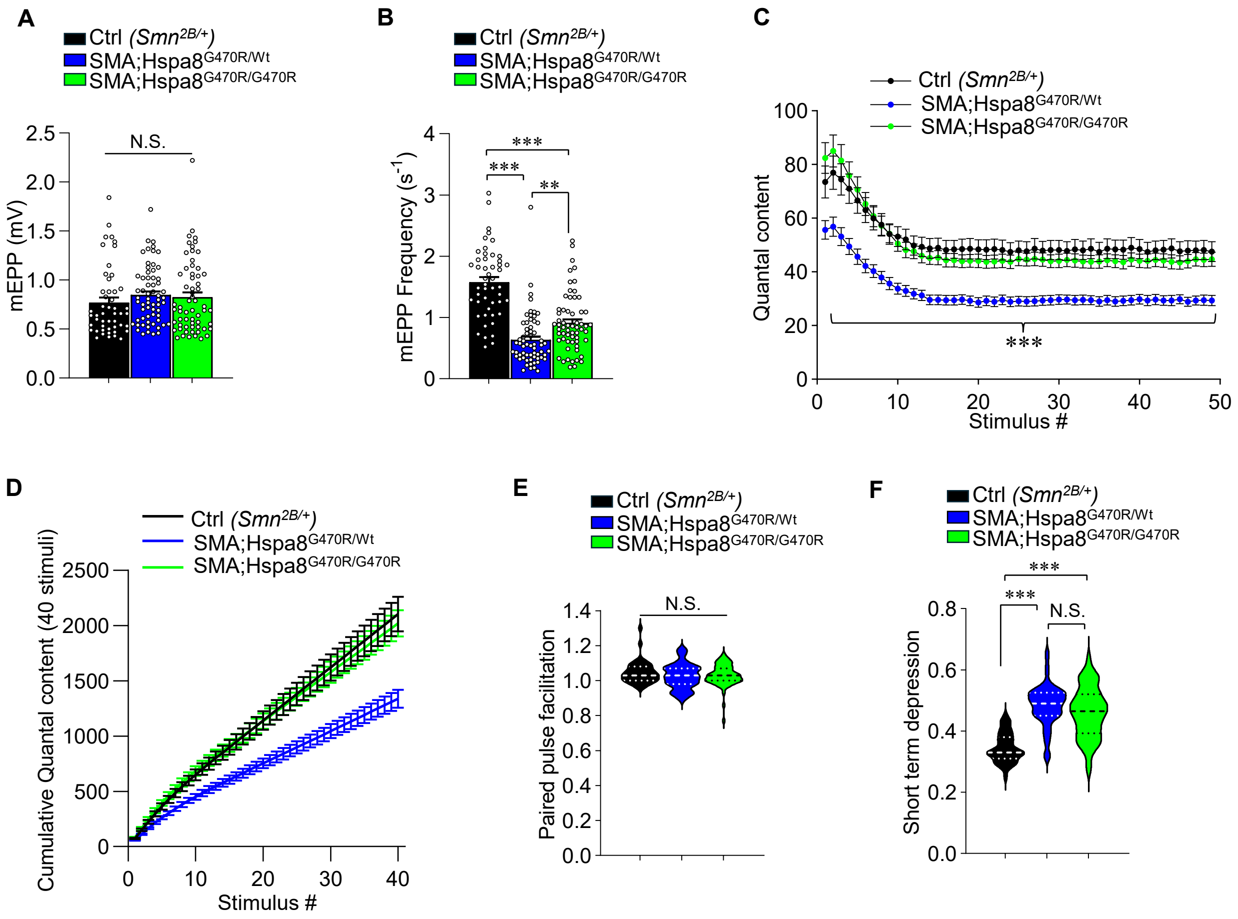


**Supplemental Figure 9 – Neurotransmission defects in young adult *$Smn^{2B/-};Hspa8^{G470R/Wt}$* mutants models type II SMA.** Quantified depictions of **(A)** mEPP amplitudes and **(B)** mEPP frequencies in 4-month-old controls and SMA mice heterozygous or homozygous for the G470R variant. Note: ***, $P < 0.001$; Kruskal-Wallis test employed for both analyses. **(C)** Quantal content in the aforementioned group of mice in response to high-frequency (20Hz, 2.5s) stimulation trains. Note: ***, $P < 0.001$, between control and *$Smn^{2B/-};Hspa8^{G470R/Wt}$* mutants, two-way ANOVA. **(D)** Representation of additive quantal content values in the three mouse cohorts as they rise with the stimulation train. Note lower trajectory of the trace corresponding to the mutant heterozygous for the modifier relative to that of the control and mice with two variant alleles. Graphs of **(E)** paired-pulse facilitation (EPP2/EPP1) and **(F)** short-term depression [1 – (mean EPP30 – 40)/EPP1] at NMJs of the young adult mice. Note: ***, $P < 0.001$, Kruskal-Wallis test (panels E, F). Also note: All analyses depicted here are based on sample sizes of n = 40 – 60 NMJs from N = 5 – 6 mice. Data: mean ± SEM
